# Role of microglia in stress-induced alcohol intake in female and male mice

**DOI:** 10.1101/2024.06.05.597614

**Authors:** Alexa R. Soares, Vernon Garcia-Rivas, Caroline Fai, Merrilee A. Thomas, Xiaoying Zheng, Marina R. Picciotto, Yann S. Mineur

## Abstract

Rates of alcohol use disorder (AUD) have escalated in recent years, with a particular increase among women. Women are more susceptible to stress-induced alcohol drinking, and preclinical data suggest that stress can increase alcohol intake in female rodents; however, a comprehensive understanding of sex-specific neurobiological substrates underlying this phenomenon is still emerging. Microglia, the resident macrophages of the brain, are essential for reshaping neuronal processes, and microglial activity contributes to overall neuronal plasticity. We investigated microglial dynamics and morphology in limbic brain structures of male and female mice following exposure to stress, alcohol or both challenges. In a modified paradigm of intermittent binge drinking (repeated “drinking in the dark”), we determined that female, but not male, mice increased their alcohol consumption after exposure to a physical stressor and re-exposure trials in the stress-paired context. Ethanol (EtOH) drinking and stress altered a number of microglial parameters, including overall number, in subregions of the amygdala and hippocampus, with effects that were somewhat more pronounced in female mice. We used the CSF1R antagonist PLX3397 to deplete microglia in female mice to determine whether microglia contribute to stress-induced escalation of EtOH intake. We observed that microglial depletion attenuated stress-induced alcohol intake with no effect in the unstressed group. These findings suggest that microglial activity can contribute to alcohol intake under stressful conditions, and highlight the importance of evaluating sex-specific mechanisms that could result in tailored interventions for AUD in women.

## Introduction

Rates of alcohol use disorder (AUD) have increased sharply among women across Western societies [1]. Despite this, women are still underrepresented in clinical studies [2], even though women are more likely to develop AUD after stress [3]. Preclinical studies have also demonstrated that stress can exacerbate alcohol consumption, particularly in female animals [4], and much research has focused on limbic structures such as the amygdala and hippocampus (HPC) [5] that are activated by stress [6,7] and alcohol [8,9]. Importantly, increased activity of the amygdala [10], and decreased hippocampal volume have been observed following chronic alcohol exposure [11], and also following exposure to chronic stress [12].

One mechanistic hypothesis to explain changes in activity and structure of limbic brain regions is the recruitment of neuroimmune signaling during initial exposure to stress or alcohol, leading to dysregulation of local signaling and neuronal plasticity that is exacerbated by adaptations in signaling following chronic exposure [13,14]. Importantly, heightened immune reactivity has consistently been observed in females [15,16], suggesting that neuroimmune signaling may contribute to sex differences in stress-related drinking behaviors [4]. Microglia are the resident macrophages of the brain [17] and are instrumental in orchestrating the brain’s response to both alcohol [18] and stress [19], with M1 macrophages predominantly contributing to pro-inflammatory reactions, while M2 macrophages primarily engage in anti-inflammatory processes [20]. Microglial activity is also important for pruning of neuronal processes and contributes to synaptic plasticity [21], although studies of sex differences in microglial function are relatively sparse [4]. Moreover, the classification of microglial activation states lacks consensus [17,22], and the multifaceted functions and phenotypes of microglia exhibit substantial regional heterogeneity [23,24].

We hypothesized that microglial function and morphology in limbic structures might be altered synergistically by exposure to stress and alcohol, and might be recruited differentially in male and female mice. We further hypothesized that differences in microglial number and function might contribute to stress-dependent drinking. We therefore optimized a behavioral paradigm of chronic alcohol drinking during which mice were exposed to a physical stressor and then placed in the stress-paired environment followed by intermittent binge trials. This paradigm resulted in a female-specific increase in alcohol drinking that was accompanied by sex- and region-specific effects on microglial number and morphology. Based on these microglial adaptations and the fact that stress increased alcohol drinking only in females, we depleted microglia pharmacologically in female mice to determine whether microglial activity might be involved in escalation of stress-induced alcohol drinking.

## Materials and Methods

### Experimental Design

#### Experiment 1: Sex differences in stress-induced EtOH drinking

Male (*n* = 24) and female (*n* = 24) C57BL6/J mice (*EtOH* group) underwent passive EtOH exposure and were then trained to acquire volitional EtOH “drinking in the dark” (DiD) in a 2-bottle choice paradigm for 11 trials (T; 5 trials/week). At the end of this phase (T11), mice were randomized into *Stress* and *No-Stress* groups (*n* = 12/sex/group). *Stress* mice were subjected to repeated foot shock prior to T12 and T13, re-exposed to the stress-paired context (without shock) on T15, T17, and T19, and finally re-exposed to foot shock on T21 and T22 (Fig. 1A). Access to EtOH was extended to a 4 hr binge period on T13 and T22. Mice were perfused upon completion of T22 and brains collected for immunohistochemistry (IHC). Control animals were age-matched, male and female mice (*n* = 24/sex) housed in the same facility but not exposed to EtOH (*No-EtOH* group). *No-EtOH* mice were also randomized into *Stress* and *No-Stress* groups (*n* = 12/sex/group), subjected to the same stress/re-exposure regimen described above, and perfused for IHC on the same day as *EtOH* mice. Full details provided in Supplemental Materials and Methods.

**Fig. 1:**
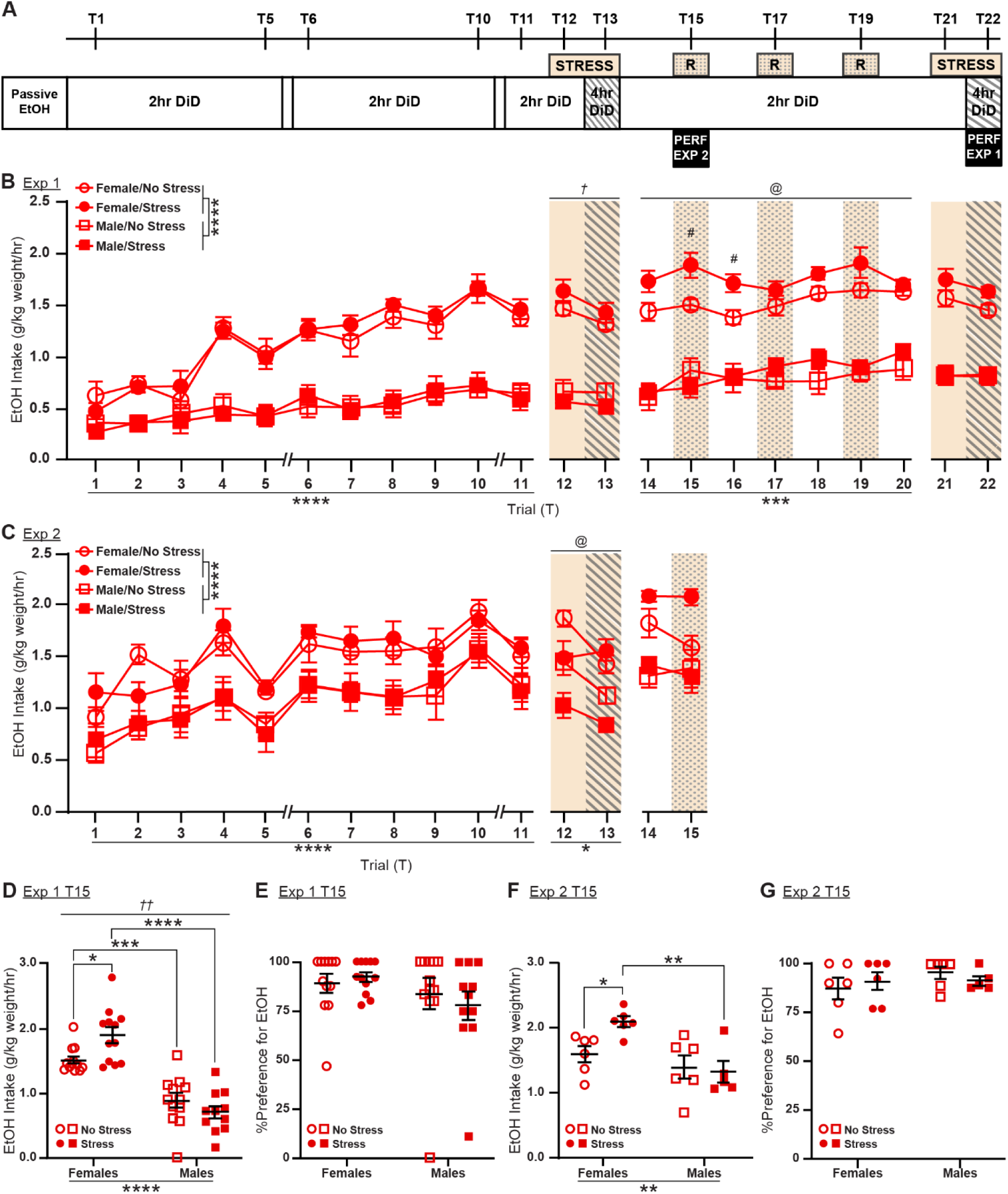
Sex differences in stress-induced EtOH drinking. **A.** <u>Experimental design of Experiment 1 (up to trial 22) and Experiment 2 (up to trial 15).</u> EtOH = ethanol, DiD = drinking in the dark; R = re-exposure to stress-paired context; PERF = perfusion. **B.** <u>Experiment 1:</u> Time course of EtOH intake across 22 trials of DiD. All mice acquired DiD over 11 training trials, with females reaching a higher intake than males. *Stress* females increased their EtOH intake during trials 14-20. *Post-hoc* analyses for these sessions in female mice revealed that the strongest stress-induced effect on EtOH intake occurred on trials 15 and 16. **C.** <u>Experiment 2</u>: Acquisition of EtOH DiD was similar to Experiment 1. **D.** <u>Trial 15 (Experiment 1):</u> *Stress* female mice showed significantly greater EtOH intake compared to *Stress* male mice and *No-Stress* female mice. **E.** There were no statistical differences in relative preference for EtOH. **F.** <u>Trial 15 (Experiment 2)</u>: *Stress* female mice showed significantly greater EtOH intake compared to *No-Stress* female mice and *Stress* male mice. **G.** There were no statistical differences in relative preference for EtOH. Exp 1: *n* = 12/group; Exp 2: *n* = 6/group. Data represent averages +/-SEM. Main effects and posthocs: ^#^*p* < 0.08; ^@,^\**p* < 0.05; \*\**p* < 0.01; \*\*\**p* < 0.001; \*\*\*\**p* < 0.0001. Sex x Stress interaction effects: ^†^*p* < 0.05; ^††^*p* < 0.01.

#### Experiment 2: Early microglial alteration following stress-induced EtOH drinking

As in Experiment 1, a separate cohort of male (*n* = 12) and female (*n* = 12) mice followed the same experimental protocol, but mice were perfused for IHC after the first re-exposure to the stress context (T15; Fig. 1A). As in Experiment 1, aged-matched *No-EtOH* mice (*n* = 6/sex/group) were subjected to the stress/re-exposure regimen described above and were perfused on the same day as *EtOH* mice.

#### Experiment 3: Role of microglia in stress-induced EtOH drinking in females

A separate cohort of female mice (*n* = 64) was passively exposed to EtOH and trained for volitional DiD for 10 Baseline (B) trials (B1-B10; Fig. S14). Mice were then randomized into a *Vehicle* diet group (*n* = 32) receiving OpenStandard Diet (Research Diets Inc) and a *PLX* diet group (*n* = 32), receiving OpenStandard Diet with PLX3397 (290 mg/kg, MedChemExpress) [25]. All mice had *ad-libitum* access to their designated chow; body weight and pellet consumption were assessed every ∼3 days. Mice continued DiD for 14 trials (T1-14), corresponding to the 21 calendar days needed for significant microglia depletion after PLX3397 chow introduction [25]. After T14, each diet group was randomized into *Stress* and *No-Stress* subgroups (*n* = 16/diet) and, starting from T15, subjected to the same stress/re-exposure regimen as in Experiment 1 (Fig. 4A).

All procedures involving animals were approved by the Yale University Committee on the Care and Use of Animals and were carried out in accordance with the Guidelines in the National Institutes of Health Guide for the Care and Use of Laboratory Animals.

### Immunohistochemistry

Following behavioral studies of EtOH and stress exposure, mice were anesthetized with pentobarbital (Fatal-Plus, Vortech Pharmaceuticals) and intracardially perfused with ∼50 ml of 4% paraformaldehyde (PFA; Electron Microscopy Sciences) in phosphate-buffered saline (PBS; Gibco) immediately after the final DiD trial. Brains were extracted and post-fixed for 1 day in 4% PFA at 4°C before transferring to 30% sucrose (Millipore Sigma) in PBS at 4°C. Brains were sectioned on a freezing microtome (Leica) at 40 μm and slices were stored in 0.02% sodium azide (Millipore Sigma) PBS solution at 4°C. Full details of IHC analyses provided in Supplemental Materials and Methods.

### Statistical analyses

All statistical analyses were conducted using GraphPad Prism 10 or SPSS (SAS software). See Supplemental Methods for full details.

## Results

### Sex differences in stress-induced EtOH drinking in mice

We first established a behavioral model that captured sex differences at the intersection of chronic EtOH intake and repeated stress exposure. Across 11 days of DiD acquisition, male and female mice progressively increased volitional EtOH drinking (Trial effect: *F*_7,327_ = 37.20; *p* < 0.0001; Fig. 1B), with a more pronounced effect in female mice (Sex effect: *F*_1,46_ = 92.90; *p* < 0.0001; Trial x Sex effect: *F*_10,450_ = 10.45; *p <* 0.0001; Fig. 1B). Relative preference for EtOH over water increased progressively during this period (Trial effect: *F*_7,301_ = 29.22; *p* < 0.0001; Fig. S2A). By the end of the acquisition phase (T11), female mice consumed on average ∼2.5x more EtOH compared to male mice (*t* = 8.807; *p* < 0.0001; Fig. 1B), but there were no sex differences in relative preference for EtOH (Fig. S2A).

Immediately following stress trials 12 and 13, EtOH intake was altered in *opposite* directions in each sex (Sex x Stress effect: *F*_1,44_ = 4.316; *p* = 0.0436, Fig. 1B), although stress-induced changes in each sex did not reach significance. In the post-stress period between T14-T20, however, a significant sex-specific increase in EtOH intake was observed among *Stress* mice (Sex x Stress x Trial effect: *F*_6,257_ = 2.731; *p* = 0.0137, Fig. 1B), with the effect of stress only observed among females (*F*_1,22_ = 12.09; *p* = 0.0021; Fig. 1B).

Despite the lack of a generalized stress effect on EtOH intake on T15, sex modulated the effect of stress during this trial (Sex x Stress effect: *F*_1,43_ = 7.784; *p* = 0.0078; Fig. 1D). Specifically, female *Stress* mice had significantly higher EtOH intake than female *No-Stress* mice (*t* = 2.736; *p* = 0.0180; Fig. 1D), but both sexes showed a strong preference for EtOH (Fig. 1E).

Measurements of blood EtOH levels were consistent with intoxication [26,27] and did not differ significantly by sex or stress exposure (Fig. S2C). Due to the high EtOH intake observed on T15, we selected this critical time point of interest for the female-specific stress-induced increase in EtOH intake for follow up in Experiment 2.

Experiment 2 largely replicated effects of Experiment 1 on acquisition of EtOH intake (T1-T11, Trial effect: *F*_5,109_ = 17.50; *p* < 0.0001; Sex effect: *F*_1,22_ = 29.35; *p* < 0.0001; Fig. 1C) and relative preference (T1-T11, Trial effect: *F*_5,118_ = 26.14; *p* < 0.0001; Sex effect: *F*_1,22_ = 19.13; *p* = 0.0002; Fig. S2B). Similarly, despite a lack of a generalized stress effect on T15, there was an interaction between sex and stress (Sex effect: *F*_1,19_ = 11.98; *p* = 0.0026; Sex x Stress interaction: *F*_1,19_ = 4.191; *p* = 0.05; Fig. 1F). Specifically, female *Stress* mice had higher EtOH intake than female *No-Stress* mice (*t* = 2.556; *p* = 0.0386; Fig. 1F), and male *Stress* mice (*t* = 3.806; *p* = 0.0024; Fig. 1F) on T15. As above, both sexes showed a strong preference for EtOH (Fig. 1G). These experiments demonstrate a reliable, female-specific escalation of EtOH drinking as a result of repeated stress exposure.

### Characterization of amygdala microglial changes following stress and EtOH exposure

We used a number of different molecular markers to determine how sex, stress exposure, and EtOH drinking affect microglial number and morphology. In Experiment 1, animals were sacrificed on T22, after the final stress exposure; in Experiment 2, animals were sacrificed on T15, the earliest time point with an observed effect of stress on drinking in Experiment 1 (Fig. 1D). We performed two triple-stain immunohistochemical analyses to measure different markers of microglial activation (Table S1). The first stain (Fig. 2A) combined expression of Iba1 (a widely-used microglial marker [28]) and P2Y12 (enriched in microglial processes [29,30]), allowing us to measure microglial density, as well as soma size, branch number, and branch length to visualize cell morphology as a readout of activation state. Smaller somas with increased branching represent a resting, ramified state; as microglia become activated, they shift towards an ameboid phenotype, with larger somas and decreased branching [31]. We also measured colocalization of CD68 with Iba1 and P2Y12 as a measure of microglial phagocytic activity [32]. In the second stain (Fig. 2B), we compared expression of iNos, an M1 marker, and Arg1, an M2 marker, in Iba1^+^ microglia; these enzymes compete for the substrate arginine, so this comparison allows for tracking of M1 (pro-inflammatory) vs M2 (anti-inflammatory) polarization [33].

**Fig. 2:**
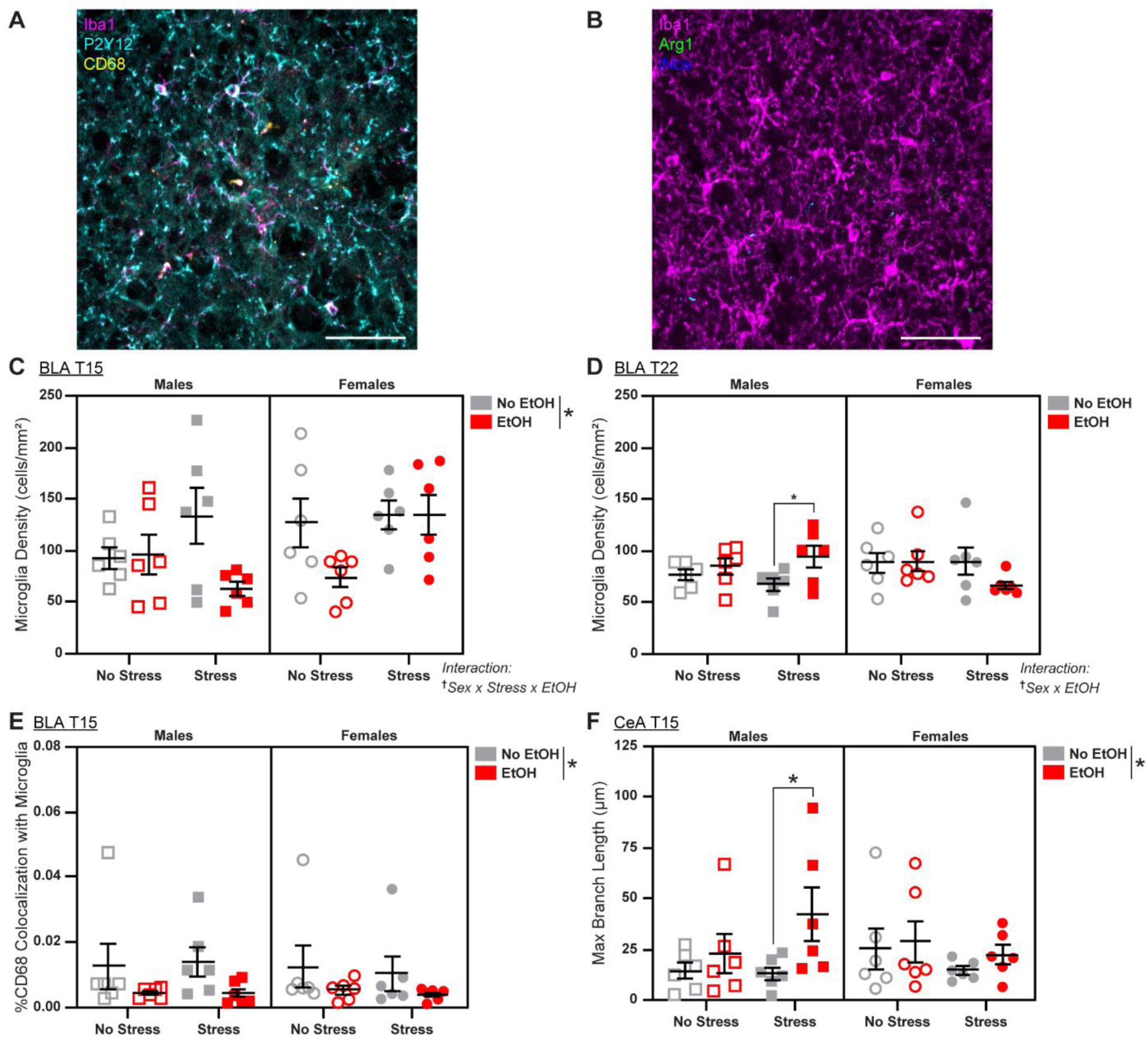
Effects of sex, stress, and EtOH on microglial phenotypes in the amygdala. **A.** Example max-intensity Z-projection for the Morphology/Phagocytosis stain. Magenta = Iba1; yellow = CD68; cyan = P2Y12; scale bar = 50 µm **B.** Example max-intensity Z-projection for the Polarization stain; Magenta = Iba1; green = Arg1; blue = iNos; scale bar = 50 µm. **C.** There is a main effect of EtOH on microglial density in the BLA on T15, which appears to be driven by reductions in males that experienced stress and females that did not. **D.** On T22, EtOH increased density in the BLA in males, particularly those that experienced stress (1 outlier excluded). **E.** EtOH decreased BLA microglial CD68 expression on T15 (1 outlier excluded). **F.** EtOH increased branch length in the CeA on T15, particularly in males that experienced stress. *n* = 5-6 animals/group. See Figs. S3 and S5 for representative micrographs and Tables S2 and S3 for detailed statistics. Data are averages +/- SEM. \**p* < 0.05; ^†^Interaction effect *p* < 0.05.

Within the amygdala, we focused on the basolateral (BLA; see Fig. S3 for representative micrographs and Table S2 for complete statistical analyses) and central (CeA; see Fig. S5 and Table S3 for complete statistical analyses) subnuclei, two regions known to regulate stress and alcohol-related behaviors [4]. EtOH altered several microglial phenotypes in these regions, particularly on T15. EtOH decreased BLA microglial density on T15 (*F*_1,40_ = 5.776; *p* = 0.0210; Fig. 2C), but stress reversed the EtOH-induced decrease in females (*F*_1,40_ = 6.728; *p* = 0.0164; Fig. 2C). By T22, a Sex x EtOH interaction was observed (*F*_1,40_ = 5.425; *p* = 0.0250; Fig. 2D); *post-hoc* analyses revealed an EtOH-induced increase in density specific to males, driven by the *Stress* group (*F*_1,20_ = 5.443; *p* = 0.0302; Table S2). EtOH also decreased CD68 expression in BLA microglia (*F*_1,40_ = 6.936; *p* = 0.0120; Fig. 2E), particularly among males (*F*_1,20_ = 4.588; *p* = 0.0447; Table S2). In the CeA, an increase in branch length (*F*_1,40_ = 4.727; *p* = 0.0357; Fig. 2F) was induced by EtOH, and *post-hoc* analyses revealed that this effect was driven by the male *Stress* group (*t* = 2.464; *p* = 0.0453; Table S3), indicating more ramified morphology in this group. These effects were limited to T15 and did not persist to T22 (Figs. S4G, S7D).

Together, these data suggest that EtOH has early anti-inflammatory effects on amygdala microglia, particularly in the male *Stress* groups, but these effects diminish with prolonged stress and EtOH exposure. Additionally, baseline sex differences were observed in both BLA and CeA, but were not consistent across timepoints (Figs. S4, S6, & S7).

### Characterization of HPC microglial changes following stress and EtOH exposure

We went on to examine microglial phenotypes in CA1 (Fig. S8, Table S4), CA3 (Fig. S10, Table S5), and dentate gyrus (DG; Fig. S12, Table S6) subfields of the HPC, revealing complex interactions between sex, stress, and EtOH. Similar to the amygdala, we saw sex differences suggesting increased baseline microglial activation in males in CA3, but these did not persist across timepoints (Fig. S11). In CA1, we observed a Sex x Stress interaction (*F*_1,40_ = 5.739; *p* = 0.0214; Fig. 3A) on T15, in which stress increased microglia soma size specifically in males (*F*_1,20_ = 4.543; *p* = 0.0456; Table S4). We also saw a Sex x Stress x EtOH interaction (*F*_1,40_ = 4.120; *p* = 0.0491; Fig. 3C) on the ratio of iNos:Arg1 levels, suggesting that stress decreases M1 polarization in females, and this is reversed by EtOH. On T22, CD68 expression revealed a Sex x EtOH interaction (*F*_1,38_ = 7.100; *p* = 0.0113; Fig. 3B) due to a male-specific EtOH-induced decrease in phagocytic activity (*F*_1,19_ = 6.971; *p* = 0.0161; Table S4) driven by the *Stress* group (*t* = 2.604; *p* = 0.0345; Table S4). In CA3, stress increased phagocytic activity on T22 (*F*_1,39_ = 6.434; *p* = 0.0153; Fig. 3F) and decreased branch number on T15 (*F*_1,39_ = 4.647; *p* = 0.0373; Fig. 3D), particularly in females (*F*_1,20_ = 6.950; *p* = 0.0158; Table S5) exposed to EtOH (*t* = 2.806; *p* = 0.0217; Table S5). Additionally, EtOH decreased microglial phagocytic activity on T15 (*p* = 0.0088; Fig. 3E). When measuring iNos/Arg1 expression in DG on T15, we saw a Stress x EtOH interaction (*F*_1,40_ = 4.784; *p* = 0.0346; Fig. 3H) and a Sex x Stress x EtOH interaction (*F*_1,40_ = 4.422; *p* = 0.0418; Fig. 3H), indicating that in males (*F*_1,20_ = 7.579; *p* = 0.0123; Table S6), stress increases M1 polarization (*t* = 2.787; *p* = 0.0266; Table S6), and this is reversed with EtOH exposure (*t* = 2.620; *p* = 0.0325; Table S6). On T22, there was a Sex x Stress x EtOH interaction (*F*_1,40_ = 4.187; *p* = 0.0473; Fig. 3G) on microglial density, because stress decreased density in females (*F*_1,20_ = 5.369; *p* = 0.0312; Table S6), particularly in the EtOH group (*t* = 2.966; *p* = 0.0152; Table S6). Together, these data reveal that stress has pro-inflammatory effects on HPC microglia in males that diminish over time and with EtOH administration. In contrast, stress appears to inhibit microglial activity and proliferation in females, and this is potentiated by EtOH in DG. The microglial IHC results from the amygdala and HPC are summarized in Table 1.

**Fig. 3:**
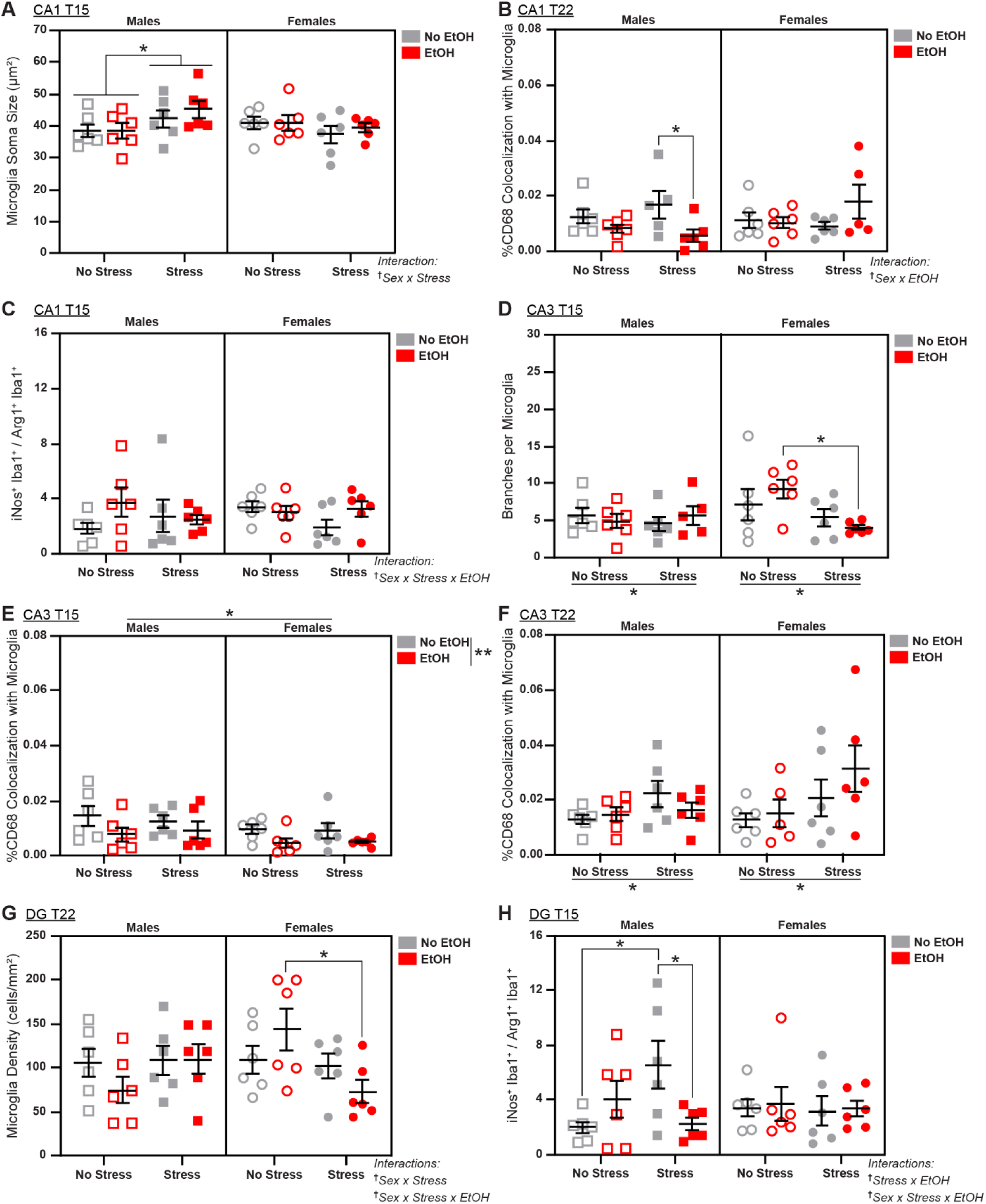
Effects of sex, stress, and EtOH on microglial phenotypes in the hippocampus. **A.** On T15, stress increased CA1 microglia soma size in males. **B.** On T22, EtOH decreased microglial CD68 expression in CA1, particularly in males that experienced stress (2 outliers excluded). **C.** In CA1 on T15, EtOH promoted M1 polarization, as indicated by iNos expression, in *No-Stress* males and *Stress* females. **D.** On T15, stress decreased microglial branch number in CA3, particularly in females (1 outlier excluded). **E.** On T15 in CA3, males showed overall increased microglial CD68 expression and EtOH decreased CD68, particularly in females. **F.** Stress increased CA3 microglial CD68 expression on T22 (1 outlier excluded). **G.** In the female DG, stress decreased microglia density, particularly in females administered EtOH. **H.** Stress promoted M1 polarization in the male DG at T15, but EtOH reversed this stress effect. *n* = 5-6 animals/group. See Figs. S8, S10, and S13 for representative micrographs and Tables S4-S6 for detailed statistics. Data are averages +/- SEM. \**p* < 0.05; ^†^Interaction effect *p* < 0.05.

**Table 1:** Summary of microglial changes following stress and EtOH exposure.

|  | **T15** | **T22** |
| --- | --- | --- |
| **Cell Density** | **EtOH** ↓ density in BLA (driven by <u>**Stress**</u> ♂ and <u>No-Stress</u> ♀) | ♂: **EtOH** ↑ density in BLA<br>♀ ↑ density in CeA |
|  |  | ♀: <u>**Stress**</u> ↓ density in DG (driven by **EtOH**) |
| **Morphology** | **EtOH** ↑ branch length in CeA |  |
|  | <u>**Stress**</u> ↓ branch number in CA3<br>♂: <u>**Stress**</u> ↑ soma size in CA1 |  |
| **Protein Markers of Activation** | **EtOH** ↓ CD68 in BLA |  |
|  | **EtOH** ↓ CD68 in CA<br>♀: <u>**Stress**</u> ↓ iNos:Arg1 ratio in CA1 (reversed by **EtOH**)<br>♂: <u>**Stress**</u> ↑ iNos:Arg1 ratio in DG (reversed by **EtOH**) | <u>**Stress**</u> ↑ CD68 in CA3<br>♂: **EtOH** ↓ CD68 in CA1 |

### Pharmacological depletion of microglia during stress and EtOH exposure

Since stress increased drinking selectively in female mice, and IHC studies revealed that stress and EtOH altered microglial density in females (Fig. 3G), we hypothesized that depleting microglia would modulate the stress-induced escalation of drinking that we observed in Experiments 1 and 2 (Figs. 1D, 1F). Over trials, female mice in Experiment 3 acquired volitional EtOH intake (*F*_4,227_ = 17.18; *p* < 0.0001; Fig. S14A) and relative preference for EtOH (*F*_5,276_ = 27.24; *p* < 0.0001; Fig. S14B) for 10 days prior to diet change, consistent with Experiments 1 and 2. Mice in the vehicle and PLX groups did not differ statistically in their EtOH drinking (Fig. S14A) or relative preference for EtOH (Fig. S14B) before diet group assignment. In the 14 trials of DiD following introduction of the vehicle or PLX diet, the new baseline EtOH intake was not significantly different between subgroups prior to stress exposure (Fig. 4B). Exposure to stress prior to trials 15 and 16 did not induce significant differences in EtOH intake between diet groups (Fig. 4B). However, EtOH intake across the following 7 days post-stress (T17-T23) was significantly increased by stress (Stress effect: *F*_1,59_ = 6.11; *p* = 0.016; Fig. 4B), with a minor effect on relative preference for EtOH (Stress effect: *F*_1,59_ = 3.30; *p* = 0.074; Fig. S14B), although there were no effects of diet on intake. Consistent with experiments 1 and 2, vehicle mice showed a stress effect on EtOH intake from T17-T23 (Stress effect: *F*_1,29_ = 7.18; *p* = 0.012; Trial effect: *F*_6,171_ = 2.175; *p* = 0.0449), but this was not observed in PLX-treated mice. *Posthoc* analyses revealed that EtOH intake in vehicle *Stress* mice was increased on T19 (*t* = 3.055, *p* = 0.016; Fig. 4B), ∼24 hrs after the first re-exposure to the stress context. On T19, vehicle *Stress* females had significantly higher EtOH intake compared to vehicle *No-Stress* (Stress x diet: *F*_1,56_ = 10.29; *p* = 0.022; Bonferroni posthoc t-test: *t* = 3.013; p = 0.0077; Fig. 4C). Despite a generalized stress effect on relative preference for EtOH on T19, there were no significant differences across diet groups (Stress effect: *F*_1,56_ = 7.144; *p* = 0.0098; Fig. S14E).

**Fig. 4:**
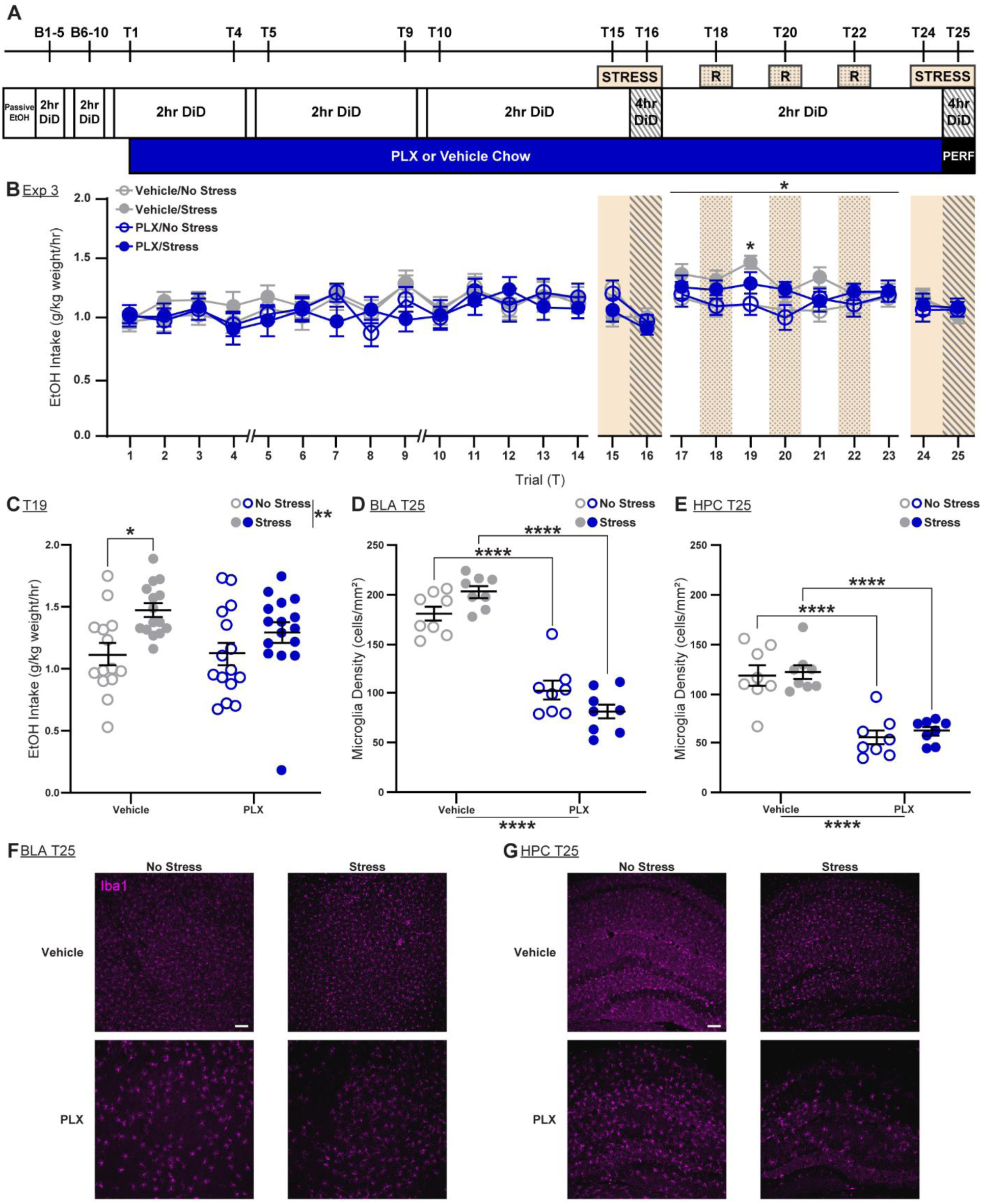
Microglial depletion with PLX prevents stress-induced EtOH drinking in female mice. **A.** <u>Experimental design for Experiment 3</u>. B1-B10 represent the acquisition of DiD prior to diet switch (see Fig. S14). EtOH = ethanol; DiD = drinking in the dark; R = re-exposure to stress-paired context; PERF = perfusion**. B.** Exposure to stress increased EtOH intake in the *Vehicle Stress* group in the period between trials 17 and 23. *n* = 8 animals/group. **C.** Trial 19: Stress increased EtOH intake. *Vehicle Stress* mice showed higher EtOH intake compared to *Vehicle No-Stress* mice, an effect diminished in the *PLX* subgroups. **D.** PLX depletes BLA microglia, in both the *Stress* and *No-Stress* groups; *n* = 8 animals/group. **E.** PLX depletes HPC microglia, in both the *Stress* and *No-Stress* groups; *n* = 8 animals/group. **F.** Representative micrographs of Iba1 staining in BLA. **G.** Representative micrographs of Iba1 staining in HPC; magenta = Iba1; scale bar = 10 μm. Data are averages +/- SEM. \**p* < 0.05; \*\**p* < 0.01; \*\*\**p* < 0.001; \*\*\*\**p* < 0.0001.

Following completion of the behavioral study, mice were euthanized and we quantified Iba1^+^ cell bodies using IHC to confirm microglial depletion in the PLX group. In both the BLA (*F*_1,28_ = 167.4; *p* < 0.0001; Fig. 4D & 4F) and HPC (*F*_1,28_ = 66.46; *p* < 0.0001; Fig. 4E & 4G) there was a dramatic decrease in microglia density in the PLX group compared to the vehicle group. Thus, microglial depletion caused by CSF1R inhibition can reverse the stress-induced increase in EtOH drinking seen in female mice.

## Discussion

Despite data suggesting that stress is more effective in potentiating alcohol drinking among women [34], only a limited number of clinical [35] and preclinical [4] studies have explored sex differences in stress-induced alcohol intake. In this study we optimized a mouse model that shows higher EtOH intake in females and a female-specific escalation of intake in response to stress exposure. This modified DiD model mimics chronic-intermittent alcohol consumption interspersed with binge episodes that is often observed in individuals with AUD [36]. In addition, volitional alcohol drinking when water or alternate rewards are available can enhance construct validity of the behavioral model [37–39].

In this study, relative preference for EtOH peaked ∼9 days after DiD training and stabilized thereafter, making this model relevant for identifying longer-term interventions that alter EtOH intake. Chronic-intermittent DiD trials resulted in stable EtOH intake even after the first stress exposure (T12-13), but we observed a robust increase in EtOH intake only in female mice after stress exposure (T15-T16). A similar delayed increase in EtOH intake in the days following inescapable foot shock exposure has been reported previously in studies of male rodents trained to drink EtOH in a 23hr two-bottle choice paradigm [40–44]; however, these studies did not include female rodents.

Recent studies have shown that repeated exposure to predator odor can promote drinking in both male and female mice [45–48], with individual variations in the proportion of animals across sexes sensitized to stress-induced intake [47,48]. The female-specific effect observed in the current study could be due to differences in the nature and timing of the stressor, as well as the schedule of EtOH accessibility. We note that the binge schedule of drinking is relevant to female-specific trends of stress-induced alcohol seeking observed in human studies [35,49–51]. We found that DiD is a reliable model for binge alcohol drinking [26,27,52,53], and results in blood EtOH levels that are consistent with intoxication similar to the human condition [26,27].

It is noteworthy that in the current behavioral paradigm, the first significant stress-induced increase in EtOH intake among female mice occurred on T15, which coincides with the first re-exposure to the stress-paired context. Foot shock stressors promote robust increases in corticosterone in rodents [54], and re-exposure to the stress context in the absence of shock can elicit strong fear- and stress-conditioned responses in rodents [55–58]. Clinical studies have shown that exposure to stress-associated cues can promote cravings for alcohol [59–61]. Our results thus suggest that a history of chronic EtOH and/or binge drinking can be exacerbated by stressful experience, particularly in female mice.

The delayed nature of this stress-induced increase in EtOH intake is likely due to long lasting cellular-, molecular- and circuit-based changes in limbic structures arising from prolonged EtOH and repeated stress exposure [62–64]. These changes could be due to altered microglial activation following exposure to stress, EtOH, and the combination. Moreover, these effects are likely influenced by sex, since inflammatory processes differ significantly between males and females [65,66].

The intersecting effects of stress and EtOH on microglia are sex-, brain region-, and time-dependent. In the amygdala, EtOH exerted a variety of anti-inflammatory effects on microglia at T15: cell density and CD68 expression were decreased in BLA, and branch length was increased in CeA. This may be surprising, as many studies have identified pro-inflammatory effects of EtOH [4], but others have observed downregulation of pro-inflammatory cytokines in the amygdala after repeated exposure to EtOH [67,68].

Interestingly, these effects did not persist at T22, and in fact, EtOH increased BLA cell density in males at the later time point, suggesting that early adaptive changes may be lost following chronic alcohol exposure. One consistent effect of EtOH was to downregulate CD68 expression, which occurred in multiple regions and timepoints, suggesting a pattern of inhibiting phagocytic activity which has been seen in other models of chronic EtOH exposure [69].

Effects of stress on microglial phenotypes were restricted to the HPC. Stress tended to have anti-inflammatory effects in females, characterized by reductions in branch number in CA3 and M1 polarization in CA1 at T15, and density in DG at T22. But this is complicated by the stress-induced increase in CD68 expression in CA3 at T22 across sexes. On the other hand, stress effects in males were more pro-inflammatory, with increased soma size in CA1 and M1 polarization in DG at T15, except for the observed decrease in branch number in CA3. Similar patterns of sex divergence in microglial responses to stress have been observed in other brain regions [70].

The effects of EtOH are antagonistic to the effect of stress in some cases, and synergistic in others. EtOH reversed stress effects on M1 polarization in both sexes, but potentiated the female-specific anti-inflammatory effect on microglia density. Repeated cycles of stress and alcohol exposure interact to drive chronic immune activation, as either can stimulate sensitization of microglia [71]; however, some research suggests that these patterns may be region-specific [72]. Overall, effects of stress and EtOH were more prominent in the HPC than the amygdala, consistent with the observation that HPC tends to display a unique vulnerability to stress and neuroinflammation, likely due to the high density of glucocorticoid receptors [73,74], which are known to mediate inflammatory responses to stress and alcohol [74,75].

Because we observed a variety of changes in microglial markers in limbic brain regions following stress and EtOH exposure, we hypothesized that microglial depletion via PLX would alter the female-specific stress-induced increase in drinking behavior. Consistent with observed changes in microglial signaling following stress and alcohol exposure, PLX prevented the stress-induced increase in EtOH drinking in female mice. It is possible that behavioral effects induced by systemic PLX exposure are driven by brain regions other than the amygdala and HPC, or by effects on circulating peripheral immune cells [76]. Notably, a study using the same PLX dose regimen found no effects on locomotion, anxiety-related behaviors, or cognitive function [25]. Previous studies testing the effects of PLX on alcohol intake have shown mixed effects on EtOH drinking; they did not include female animals, nor did they test effects of stress [77,78]. In the current study, PLX prevented stress-induced binge drinking in females, highlighting a potential role for regulation of microglia as a treatment for AUD in women, and particularly in the intersection between stress and problematic drinking.

Clinical trials have explored the safety and efficacy of PLX in treating various forms of cancer and found it to be well tolerated [79,80]. The current study suggests that PLX or other microglial inhibitors might be useful in treatment of AUD and suggests a novel avenue for medication development that might be particularly efficacious for women with AUD and co-morbid stress disorders.

## Acknowledgments

We would like to thank Drs. Kelsea Gildawie, Staci Bilbo, and Evan Bordt for guidance in developing the IHC protocols, Samantha Sheppard for assistance with animal husbandry, and Nadia Jordan-Spasov for assistance with reagents.

## Contributions

ARS designed, carried out, and analyzed IHC experiments, contributed to behavioral experiments, and wrote the manuscript. VGR designed, carried out, and analyzed behavioral experiments, and wrote the manuscript. MAT designed, carried out, and analyzed behavioral experiments. CF and XZ contributed to IHC analyses. MRP and YSM secured funding, gathered pilot data, designed the study, contributed to data analyses and edited the manuscript. All authors reviewed and approved the manuscript.

## References

1. Grant BF, Chou SP, Saha TD, Pickering RP, Kerridge BT, Ruan WJ, et al. Prevalence of 12-Month Alcohol Use, High-Risk Drinking, and DSM-IV Alcohol Use Disorder in the United States, 2001-2002 to 2012-2013: Results From the National Epidemiologic Survey on Alcohol and Related Conditions. JAMA Psychiatry. 2017;74:911.

2. Verplaetse TL, Cosgrove KP, Tanabe J, McKee SA. Sex/gender differences in brain function and structure in alcohol use: a narrative review of neuroimaging findings over the last 10 years. J Neurosci Res. 2021;99:309.

3. Verplaetse TL, Moore KE, Pittman BP, Roberts W, Oberleitner LM, Smith PH, et al. Intersection of Stress and Gender in Association With Transitions in Past Year DSM-5 Substance Use Disorder Diagnoses in the United States. Chronic Stress. 2018;2.

4. Mineur YS, Garcia-Rivas V, Thomas MA, Soares AR, McKee SA, Picciotto MR. Sex differences in stress-induced alcohol intake: a review of preclinical studies focused on amygdala and inflammatory pathways. Psychopharmacology (Berl). 2022;239:2041– 2061.

5. Koob GF, Volkow ND. Neurobiology of addiction: a neurocircuitry analysis. Lancet Psychiatry. 2016;3:760–773.

6. Nasrullah N, Khorashad Sorouri B, Lundmark A, Seiger R, Savic I. Occupational stress is associated with sex and subregion specific modifications of the amygdala volumes. Stress. 2023;26:2247102.

7. Olave FA, Aguayo FI, Román-Albasini L, Corrales WA, Silva JP, González PI, et al. Chronic restraint stress produces sex-specific behavioral and molecular outcomes in the dorsal and ventral rat hippocampus. Neurobiol Stress. 2022;17:100440.

8. Logrip ML, Oleata C, Roberto M. Sex differences in responses of the basolateral-central amygdala circuit to alcohol, corticosterone and their interaction. Neuropharmacology. 2017;114:123–134.

9. Maynard ME, Barton EA, Robinson C, Wooden J, Leasure JL. Sex Differences in Hippocampal Damage, Cognitive Impairment, and Trophic Factor Expression in an Animal Model of an Alcohol Use Disorder. Brain Struct Funct. 2018;223:195–210.

10. Gorka SM, Fitzgerald DA, King AC, Phan KL. Alcohol attenuates amygdala– frontal connectivity during processing social signals in heavy social drinkers. Psychopharmacology (Berl). 2013;229:141–154.

11. Wilson S, Bair JL, Thomas KM, Iacono WG. Problematic alcohol use and reduced hippocampal volume: a meta-analytic review. Psychol Med. 2017;47:2288– 2301.

12. Lindgren L, Bergdahl J, Nyberg L. Longitudinal Evidence for Smaller Hippocampus Volume as a Vulnerability Factor for Perceived Stress. Cereb Cortex. 2016;26:3527–3533.

13. Corrigan M, O’Rourke AM, Moran B, Fletcher JM, Harkin A. Inflammation in the pathogenesis of depression: a disorder of neuroimmune origin. Neuronal Signal. 2023;7:NS20220054.

14. Pascual M, Montesinos J, Guerri C. Role of the innate immune system in the neuropathological consequences induced by adolescent binge drinking. J Neurosci Res. 2018;96:765–780.

15. Migliore L, Nicolì V, Stoccoro A. Gender Specific Differences in Disease Susceptibility: The Role of Epigenetics. Biomedicines. 2021;9:652.

16. Hayter SM, Cook MC. Updated assessment of the prevalence, spectrum and case definition of autoimmune disease. Autoimmun Rev. 2012;11:754–765.

17. Woodburn SC, Bollinger JL, Wohleb ES. The semantics of microglia activation: neuroinflammation, homeostasis, and stress. J Neuroinflammation. 2021;18:258.

18. Henriques JF, Portugal CC, Canedo T, Relvas JB, Summavielle T, Socodato R. Microglia and alcohol meet at the crossroads: Microglia as critical modulators of alcohol neurotoxicity. Toxicol Lett. 2018;283:21–31.

19. Frank MG, Fonken LK, Watkins LR, Maier SF. Microglia: neuroimmune-sensors of stress. Semin Cell Dev Biol. 2019;94:176–185.

20. Yunna C, Mengru H, Lei W, Weidong C. Macrophage M1/M2 polarization. Eur J Pharmacol. 2020;877:173090.

21. Guedes JR, Ferreira PA, Costa JM, Cardoso AL, Peça J. Microglia-dependent remodeling of neuronal circuits. J Neurochem. 2022;163:74–93.

22. Ransohoff RM. A polarizing question: do M1 and M2 microglia exist? Nat Neurosci. 2016;19:987–991.

23. Masuda T, Sankowski R, Staszewski O, Prinz M. Microglia Heterogeneity in the Single-Cell Era. Cell Rep. 2020;30:1271–1281.

24. Lawson LJ, Perry VH, Dri P, Gordon S. Heterogeneity in the distribution and morphology of microglia in the normal adult mouse brain. Neuroscience. 1990;39:151– 170.

25. Elmore MRP, Najafi AR, Koike MA, Dagher NN, Spangenberg EE, Rice RA, et al. CSF1 receptor signaling is necessary for microglia viability, which unmasks a cell that rapidly repopulates the microglia-depleted adult brain. Neuron. 2014;82:380–397.

26. Rhodes JS, Ford MM, Yu C-H, Brown LL, Finn DA, Garland Jr T, et al. Mouse inbred strain differences in ethanol drinking to intoxication. Genes Brain Behav. 2007;6:1–18.

27. Thiele TE, Crabbe JC, Boehm II SL. “Drinking in the Dark” (DID): A Simple Mouse Model of Binge-Like Alcohol Intake. Curr Protoc Neurosci. 2014;68:9.49.1-9.49.12.

28. Jurga AM, Paleczna M, Kuter KZ. Overview of General and Discriminating Markers of Differential Microglia Phenotypes. Front Cell Neurosci. 2020;14.

29. Sipe GO, Lowery, RL, Tremblay M-È, Kelly EA, Lamantia CE, Majewska AK. Microglial P2Y12 is necessary for synaptic plasticity in mouse visual cortex. Nat Commun. 2016;7:10905.

30. Haynes PR, Christmann BL, Griffith LC. A single pair of neurons links sleep to memory consolidation in Drosophila melanogaster. eLife. 2015;4:e03868.

31. Lier J, Streit WJ, Bechmann I. Beyond Activation: Characterizing Microglial Functional Phenotypes. Cells. 2021;10:2236.

32. Walker DG, Lue L-F. Immune phenotypes of microglia in human neurodegenerative disease: challenges to detecting microglial polarization in human brains. Alzheimers Res Ther. 2015;7.

33. Cherry JD, Olschowka JA, O’Banion MK. Neuroinflammation and M2 microglia: the good, the bad, and the inflamed. J Neuroinflammation. 2014;11:98.

34. Curlee J. A Comparison of Male and Female Patients at an Alcoholism Treatment Center. J Psychol. 1970;74:239–247.

35. Peltier MR, Verplaetse TL, Mineur YS, Petrakis IL, Cosgrove KP, Picciotto MR, et al. Sex differences in stress-related alcohol use. Neurobiol Stress. 2019;10.

36. Bobashev GV, Liao D, Hampton J, Helzer JE. Individual patterns of alcohol use. Addict Behav. 2014;39:934–940.

37. Ahmed SH. Trying to make sense of rodents’ drug choice behavior. Prog Neuropsychopharmacol Biol Psychiatry. 2018;87:3–10.

38. Augier E, Barbier E, Dulman RS, Licheri V, Augier G, Domi E, et al. A molecular mechanism for choosing alcohol over an alternative reward. Science. 2018;360:1321– 1326.

39. Cannella N, Ubaldi M, Masi A, Bramucci M, Roberto M, Bifone A, et al. Building better strategies to develop new medications in Alcohol Use Disorder: Learning from past success and failure to shape a brighter future. Neurosci Biobehav Rev. 2019;103:384–398.

40. Matthews DB, Morrow AL, O’Buckley T, Flanigan TJ, Berry RB, Cook MN, et al. Acute mild footshock alters ethanol drinking and plasma corticosterone levels in C57BL/6J male mice, but not DBA/2J or A/J male mice. Alcohol. 2008;42:469–476.

41. Anisman H, Waller TG. Effects of inescapable shock and shock-produced conflict on self selection of alcohol in rats. Pharmacol Biochem Behav. 1974;2:27–33.

42. Racz I, Bilkei-Gorzo A, Toth ZE, Michel K, Palkovits M, Zimmer A. A Critical Role for the Cannabinoid CB1 Receptors in Alcohol Dependence and Stress-Stimulated Ethanol Drinking. J Neurosci. 2003;23:2453–2458.

43. Volpicelli JR. Uncontrollable Events and Alcohol Drinking. Br J Addict. 1987;82:381–392.

44. Volpicelli JR, Ulm RR, Hopson N. The Bidirectional Effects of Shock on Alcohol Preference in Rats. Alcohol Clin Exp Res. 1990;14:913–916.

45. Finn DA, Helms ML, Nipper MA, Cohen A, Jensen JP, Devaud LL. Sex differences in the synergistic effect of prior binge drinking and traumatic stress on subsequent ethanol intake and neurochemical responses in adult C57BL/6J mice. Alcohol. 2018;71:33–45.

46. Cozzoli DK, Tanchuck-Nipper MA, Kaufman MN, Horowitz CB, Finn DA. Environmental stressors influence limited-access ethanol consumption by C57BL/6J mice in a sex-dependent manner. Alcohol. 2014;48:741–754.

47. Alavi M, Ryabinin AE, Helms ML, Nipper MA, Devaud LL, Finn DA. Sensitivity and Resilience to Predator Stress-Enhanced Ethanol Drinking Is Associated With Sex-Dependent Differences in Stress-Regulating Systems. Front Behav Neurosci. 2022;16.

48. Nipper MA, Helms ML, Finn DA, Ryabinin AE. Stress-enhanced ethanol drinking does not increase sensitivity to the effects of a CRF-R1 antagonist on ethanol intake in male and female mice. Alcohol. 2024. 5 January 2024. 10.1016/j.alcohol.2024.01.001.

49. Breslow RA, Castle I-JP, Chen CM, Graubard BI. Trends in Alcohol Consumption Among Older Americans: National Health Interview Surveys, 1997 to 2014. Alcohol Clin Exp Res. 2017;41:976–986.

50. Chou K-L, Liang K, Mackenzie CS. Binge Drinking and Axis I Psychiatric Disorders in Community-Dwelling Middle-Aged and Older Adults: Results From the National Epidemiologic Survey on Alcohol and Related Conditions (NESARC). J Clin Psychiatry. 2011;72:1895.

51. Patock-Peckham JA, Corbin WR, Smyth H, Canning JR, Ruof A, Williams J. Effects of stress, alcohol prime dose, and sex on ad libitum drinking. Psychol Addict Behav. 2022;36:871–884.

52. Rhodes JS, Best K, Belknap JK, Finn DA, Crabbe JC. Evaluation of a simple model of ethanol drinking to intoxication in C57BL/6J mice. Physiol Behav. 2005;84:53– 63.

53. Blednov YA, Ponomarev I, Geil C, Bergeson S, Koob GF, Harris RA. Neuroimmune regulation of alcohol consumption: behavioral validation of genes obtained from genomic studies. Addict Biol. 2012;17:108–120.

54. Shanks N, Griffiths J, Zalcman S, Zacharko RM, Anisman H. Mouse strain differences in plasma corticosterone following uncontrollable footshock. Pharmacol Biochem Behav. 1990;36:515–519.

55. Li S, Murakami Y, Wang M, Maeda K, Matsumoto K. The effects of chronic valproate and diazepam in a mouse model of posttraumatic stress disorder. Pharmacol Biochem Behav. 2006;85:324–331.

56. Maier SF. Exposure to the stressor environment prevents the temporal dissipation of behavioral depression/learned helplessness. Biol Psychiatry. 2001;49:763–773.

57. Pynoos RS, Ritzmann RF, Steinberg AM, Goenjian A, Prisecaru I. A behavioral animal model of posttraumatic stress disorder featuring repeated exposure to situational reminders. Biol Psychiatry. 1996;39:129–134.

58. Sanford LD, Yang L, Wellman LL, Liu X, Tang X. Differential Effects of Controllable and Uncontrollable Footshock Stress on Sleep in Mice. Sleep. 2010;33:621–630.

59. Blaine SK, Nautiyal N, Hart R, Guarnaccia J b., Sinha R. Craving, cortisol and behavioral alcohol motivation responses to stress and alcohol cue contexts and discrete cues in binge and non-binge drinkers. Addict Biol. 2019;24:1096–1108.

60. Sinha R, Fox HC, Hong KA, Bergquist K, Bhagwagar Z, Siedlarz KM. Enhanced Negative Emotion and Alcohol Craving, and Altered Physiological Responses Following Stress and Cue Exposure in Alcohol Dependent Individuals. Neuropsychopharmacology. 2009;34:1198–1208.

61. Sinha R. How Does Stress Lead to Risk of Alcohol Relapse? Alcohol Res Curr Rev. 2012;34:432–440.

62. Becker HC, Lopez MF, Doremus-Fitzwater TL. Effects of stress on alcohol drinking: a review of animal studies. Psychopharmacology (Berl). 2011;218:131–156.

63. Noori HR, Helinski S, Spanagel R. Cluster and meta-analyses on factors influencing stress-induced alcohol drinking and relapse in rodents. Addict Biol. 2014;19:225–232.

64. Volpicelli J, Balaraman G, Hahn J, Wallace H, Bux D. The Role of Uncontrollable Trauma in the Development of PTSD and Alcohol Addiction. Alcohol Res Health. 1999;23:256–262.

65. Cyr B, de Rivero Vaccari JP. Sex Differences in the Inflammatory Profile in the Brain of Young and Aged Mice. Cells. 2023;12:1372.

66. Martínez de Toda I, González-Sánchez M, Díaz-Del Cerro E, Valera G, Carracedo J, Guerra-Pérez N. Sex differences in markers of oxidation and inflammation. Implications for ageing. Mech Ageing Dev. 2023;211:111797.

67. Doremus-Fitzwater TL, Buck HM, Bordner KA, Richey L, Jones ME, Deak T. Intoxication- and withdrawal-dependent expression of central and peripheral cytokines following initial ethanol exposure. Alcohol Clin Exp Res. 2014;38:2186–2198.

68. Gano A, Doremus-Fitzwater TL, Deak T. Sustained alterations in neuroimmune gene expression after daily, but not intermittent, alcohol exposure. Brain Res. 2016;1646:62–72.

69. Lowe PP, Morel C, Ambade A, Iracheta-Vellve A, Kwiatkowski E, Satishchandran A, et al. Chronic alcohol-induced neuroinflammation involves CCR2/5-dependent peripheral macrophage infiltration and microglia alterations. J Neuroinflammation. 2020;17:296.

70. Bollinger JL. Uncovering microglial pathways driving sex-specific neurobiological effects in stress and depression. Brain Behav Immun - Health. 2021;16:100320.

71. Crews FT, Lawrimore CJ, Walter TJ, Coleman LG. The role of neuroimmune signaling in alcoholism. Neuropharmacology. 2017;122:56–73.

72. Walter TJ, Vetreno RP, Crews FT. Alcohol and Stress Activation of Microglia and Neurons: Brain Regional Effects. Alcohol Clin Exp Res. 2017;41:2066–2081.

73. Gulyaeva NV. Stress-Associated Molecular and Cellular Hippocampal Mechanisms Common for Epilepsy and Comorbid Depressive Disorders. Biochem Mosc. 2021;86:641–656.

74. Bolshakov AP, Tret’yakova LV, Kvichansky AA, Gulyaeva NV. Glucocorticoids: Dr. Jekyll and Mr. Hyde of Hippocampal Neuroinflammation. Biochem Mosc. 2021;86:156–167.

75. Finn DA. Stress and gonadal steroid influences on alcohol drinking and withdrawal, with focus on animal models in females. Front Neuroendocrinol. 2023;71:101094.

76. Basilico B, Ferrucci L, Khan A, Di Angelantonio S, Ragozzino D, Reverte I. What microglia depletion approaches tell us about the role of microglia on synaptic function and behavior. Front Cell Neurosci. 2022;16:1022431.

77. Warden AS, Triplett TA, Lyu A, Grantham EK, Azzam MM, DaCosta A, et al. Microglia depletion and alcohol: Transcriptome and behavioral profiles. Addict Biol. 2021;26:e12889.

78. Warden AS, Wolfe SA, Khom S, Varodayan FP, Patel RR, Steinman MQ, et al. Microglia control escalation of drinking in alcohol dependent mice: Genomic and synaptic drivers. Biol Psychiatry. 2020;88:910–921.

79. Moskowitz CH, Younes A, de Vos S, Bociek RG, Gordon LI, Witzig TE, et al. CSF1R Inhibition by PLX3397 in Patients with Relapsed or Refractory Hodgkin Lymphoma: Results From a Phase 2 Single Agent Clinical Trial. Blood. 2012;120:1638.

80. Butowski N, Colman H, De Groot JF, Omuro AM, Nayak L, Wen PY, et al. Orally administered colony stimulating factor 1 receptor inhibitor PLX3397 in recurrent glioblastoma: an Ivy Foundation Early Phase Clinical Trials Consortium phase II study. Neuro-Oncol. 2015;18:557–564.

