## Supplemental Files for "Role of microglia in stress-induced alcohol intake in female and male mice"

**Supplemental Materials and Methods**

Mice

Male and female (defined by large vs. small anogenital distance at weaning) C57BL/6J mice of 8-10 weeks of age (The Jackson Laboratory) were maintained on a reverse 12 hr light/dark cycle (lights off at 11:00 AM) with *ad libitum* access to food and water. For 1 week of acclimation, mice were group-housed and handled daily, before switching to single-housing for the remainder of the experiment. All procedures were approved by the Yale University Institutional Animal Care and Use Committee in compliance with the National Institute of Health's Guide for the Care and Use of Laboratory Animals.

Sippers

Sipper tubes for delivery of alcohol were constructed by fitting a Hydropac® valve (Avidity Science) into the cut end of a graduated 5 mL syringe and fixing the valve with a plastic adhesive. Custom 3D-printed holders were custom-designed to hold two drinking sippers in the home cage (inspired by [1]). Sippers were checked daily for liquid leakage prior to drinking trials and replaced accordingly.

Passive EtOH Exposure

To acclimate mice to the taste and pharmacological effects of EtOH before 2-bottle volitional choice of EtOH vs. water, mice were exposed to increasing concentrations of alcohol diluted in water (5% EtOH v/v for 3 days, 10% EtOH v/v for 4 days) as their sole liquid source for 1 week. Bottles were weighed every 3-4 days to measure intake. On the last day of passive EtOH exposure, bottles were replaced with regular water bottles prior to lights off. EtOH solutions were prepared fresh by diluting undenatured 100% EtOH (Decon Laboratories) in filtered drinking water.

Drinking in the Dark

Mice were trained to drink EtOH in their home cage through a limited access paradigm (Drinking-in-the-Dark: DiD), based on [2] and access to EtOH was limited to 5 days a week. 3 hrs after lights off, water bottles were removed and two sippers, one containing water and one containing EtOH (10% v/v diluted in water) were introduced into the home cage for 2 hrs. Fluid volumes were measured to calculate daily EtOH intake and relative preference of EtOH and water. To prevent side bias, EtOH and water sippers were swapped every day. On selected days noted above, access to EtOH was extended to 4 hrs to mimic a drinking binge to escalate intake.

Stress exposure regimen

On stress exposure days, *Stress* mice were placed individually into a chamber containing an electrified floor grate (Med Associates, VT) inside a sound-attenuating box. Mice received 120 low-intensity inescapable footshocks (0.3 mA, 4 sec), delivered at semi-random intervals (1-17 sec) over the course of 1 hr [3–6]. A white light was turned on at the time of the first foot shock and remained on until the stress trial ended. On stress re-exposure days, *Stress* mice were placed into the stress-associated chamber for 5 min with the white light on, but received no footshocks. Stress and re-exposure trials occurred towards the end of the light phase, at least 4 hr prior to the DiD trial. *No-Stress* mice remained in their home cages undisturbed.

Measurement of blood alcohol levels

A separate group of Male (*n* = 11) and female (*n* = 11) C57BL6/J mice underwent the same EtOH and stress protocol. Immediately after completion of the last 4 hr binge session, mice were euthanized, trunk blood collected in MiniCollect® tubes containing lithium heparin (Greiner Bio-One), and immediately placed on ice. Blood samples were spun at 2000 RCF for 10 minutes at 4ºC and the resulting plasma aliquoted in 30 µl fractions, which were immediately frozen at -80ºC until assayed. Blood EtOH concentrations were measured using the EnzyChrom™ EtOH Colorimetric Assay Kit (BioAssay Systems, ECET-100) according to manufacturer's instructions.

Immunohistochemistry (IHC)

For IHC analyses, sections were washed in PBS and incubated in a 0.3% Triton-X (American Bioanalytical) PBS solution for 15 min at room temperature (RT). Sections were then rinsed in PBS for 5 min and incubated in 0.01 M citric acid (Millipore Sigma) for 30 min at 70°C for antigen retrieval, followed by PBS rinsing for 5 min and then 1 hr in a sodium tetraborate (Millipore Sigma) buffer at RT. After subsequent PBS rinsing for 5 min, sections were blocked in 3% normal donkey serum (NDS; Jackson Immuno) and 0.3% Triton-X in PBS for 1 hr at RT. Sections were then incubated in primary antibodies (Table S1) and PBS overnight at 4°C. On day 2, sections were rinsed in PBS for 5 min and incubated in secondary antibodies (Table S1) in PBS for 1 hr at RT. Sections were rinsed in PBS for 5 min, then mounted on slides and coverslipped with ProLong Gold antifade mounting medium (Invitrogen).

For Experiments 1 and 2, Z-stack images were acquired in 5 serial sections at 60x magnification with a 1.0 µm increment between slices (Fig. S1A) using a FLUOVIEW FB10i confocal microscope (Olympus). Acquired images were analyzed in FIJI [7] and Matlab using custom codes to measure microglial density, morphology (Fig. S1), phagocytic activity, and polarization state. For Experiment 3, multichannel images were acquired in 5 serial sections (spaced by 240 µm) at 10x magnification. Acquired images were analyzed in FIJI [7] and Matlab to quantify microglial density.

<u>Morphology Analyses</u>

Multichannel Z-stack images were acquired in 5 serial sections at 60x magnification using a FLUOVIEW FV10i confocal microscope (Olympus). Regions of interest were outlined using pre-defined boundaries and landmarks in accordance with a mouse brain atlas [8]. 2-5 images were captured within the BLA and CeA. 2-3 images were captured for each HPC subfield. Acquired images were analyzed first in FIJI [7] using a custom Macro code based on Young & Morrison [9] and subsequently in Matlab using custom code. Green (Iba1) and blue (P2Y12) channels were merged to form a single representation of microglia. A Z-Projection of this *Composite stack* was created based on *Max Intensity* and converted to *Grayscale. Unsharp Mask* and *Despeckle* were used to sharpen edges and remove noise. Each soma was manually counted using the Multipoint tool. Microglial density was determined by summing this count across each image for a given region of interest and dividing by the total area imaged for that region in each animal. Soma size was determined by tracing each soma using the *Freehand tool* and calculating the area using the *Measure* tool, then dividing by the total number of somas for each region of interest in each animal. The de-noised *Composite Z-projection* was processed using the *Skeletonize* tool and then the *Skeleton* plugin to quantify branch number and branch, both of which were normalized to the total number of somas.

<u>Phagocytosis Analysis</u>

Multichannel Z-stacks were acquired as described above. Acquired images were analyzed first in FIJI [7] using a custom Macro code and subsequently in Matlab using custom code. Green (Iba1) and blue (P2Y12) channels were merged to form a single representation of the microglia. The *Composite* stack was converted to 8-bit and then background was corrected across the stack using the *Threshold* tool before binarizing and *Despeckling* to remove noise. The *Analyze Particles* tool was used to quantify the total area covered by microglia as represented by expression of Iba1 and/or P2Y12, and this number was summed across all the images for a given region of interest for each animal. The stack for the red channel was similarly Thresholded, binarized, and Despeckled. The *Image Calculator* tool was used to calculate colocalization between the red stack and the Composite stack, and this was summed across images and normalized to the total Iba1/P2Y12$^+$ area to determine the percentage of CD68 expression in microglia.

Polarization Analysis

Multichannel Z-stacks were acquired as described above. Acquired images were analyzed first in FIJI [7] using a custom Macro code and subsequently in Matlab using custom code. Channels were split. The blue (Iba1) channel was duplicated and a Z-Projection was created using Max Intensity to measure microglial density as described above. To measure colocalization, the original blue stack was used. Background was corrected across the stack using the *Threshold* tool before binarizing and *Despeckling* to remove noise. The *Analyze Particles* tool was used to measure the Iba1$^+$ area. The *Image Calculator* tool was used to calculate colocalization between the red (iNos) and blue channels, and this was summed across all of the images for a given region of interest and normalized to the total Iba1$^+$ area to determine percentage of iNos expression in microglia for each animal. These steps were repeated using the green (Arg1) and blue channels to determine the percentage of Arg1 expression in microglia. The percentage of iNos colocalization was divided by the percentage of Arg1 colocalization to determine the ratio of iNos:Arg1 expression in microglia.

Depletion Analysis

Multichannel images were acquired in 5 serial sections at 10x magnification. Regions of interest were outlined using pre-defined boundaries and landmarks in accordance with a mouse brain atlas [8]. 2-3 images were captured for BLA and for the entire HPC. Acquired images were analyzed first in FIJI [7] using a custom Macro code and subsequently in Matlab using custom code. The *Polygon* tool was used to limit the region of interest to the boundaries of the BLA or HPC and the *Measure* tool was used to calculate the area, which was summed across all of the images for a given region of interest in each animal. Channels were split and the blue (Iba1) channel was used. Microglial density was determined by manually counting each soma using the *Multipoint* tool; this number was summed across all of the images for a given region of interest and divided by the total area imaged within the region of interest for each animal.

Statistical analyses

Results are presented as the mean ± standard error of the mean (SEM) and “Trial (T)” represents days when mice were exposed to EtOH. For all behavioral data, repeated-measures Analysis of Variance (ANOVA) were performed with Trial as a within-subject factor and Sex, Stress, and EtOH as between-subject factors. Because the behavioral paradigm comprised several phases with different treatments (i.e. phase 1 with exposure to daily EtOH/DiD, followed by phases with the introduction of stressors), data were analyzed independently between phases. Bonferroni tests were performed *post-hoc* for further investigation of multiple comparisons when appropriate. In the case of any sipper leakage during a drinking trial (as determined by an empty sipper tube at the end of a trial, whether EtOH or water), the drinking data from that mouse for that trial was recorded as “leaked” and removed from the final statistical analysis, and data were analyzed in a mixed effects model. For IHC data for Experiments 1 and 2, 3-way ANOVAs were performed with Sex, Stress, and EtOH as between-subject factors. Significant main effects or interactions were followed up with 2-way ANOVAs within Sex (Stress x EtOH) and post-hoc Šídák's multiple comparisons tests when appropriate. 2-way ANOVAs were used to analyze the data for Experiment 3 with Stress and Diet as between-subject factors and *post-hoc* Šídák's multiple comparisons tests when appropriate. Outliers were determined using Grubbs’ test and subsequently excluded.

### Supplementary Tables and Figures

Table S1: Antibody information for IHC experiments

| Analysis | [Concentration] Primary Antibody (Source) | [Concentration] Secondary Antibody (Source) |
|---|---|---|
| Morphology/Phagocytosis | [1:1000] rabbit anti-P2Y12 (AnaSpec) | [1:2000] donkey anti-rabbit Alexa 647 (Invitrogen) |
| Morphology/Phagocytosis | [1:1000] goat anti-Iba1 (WAKO) | [1:1000] donkey anti-goat Alexa 488 (Invitrogen) |
| Morphology/Phagocytosis | [1:1000] rat anti-CD68 (BioRad) | [1:1000] donkey anti-rat Alexa 555 (Abcam) |
| Polarization/Depletion | [1:1000] rabbit anti-iNos (Abcam) | [1:2000] donkey anti-rabbit Alexa 555 (Invitrogen) |
| Polarization/Depletion | [1:1000] goat anti-Iba1 (WAKO) | [1:1000] donkey anti-goat Alexa 647 (Invitrogen) |
| Polarization/Depletion | [1:500] chicken anti-Arg1 | [1:1000] donkey anti-chicken Alexa 488 (Millipore Sigma) |

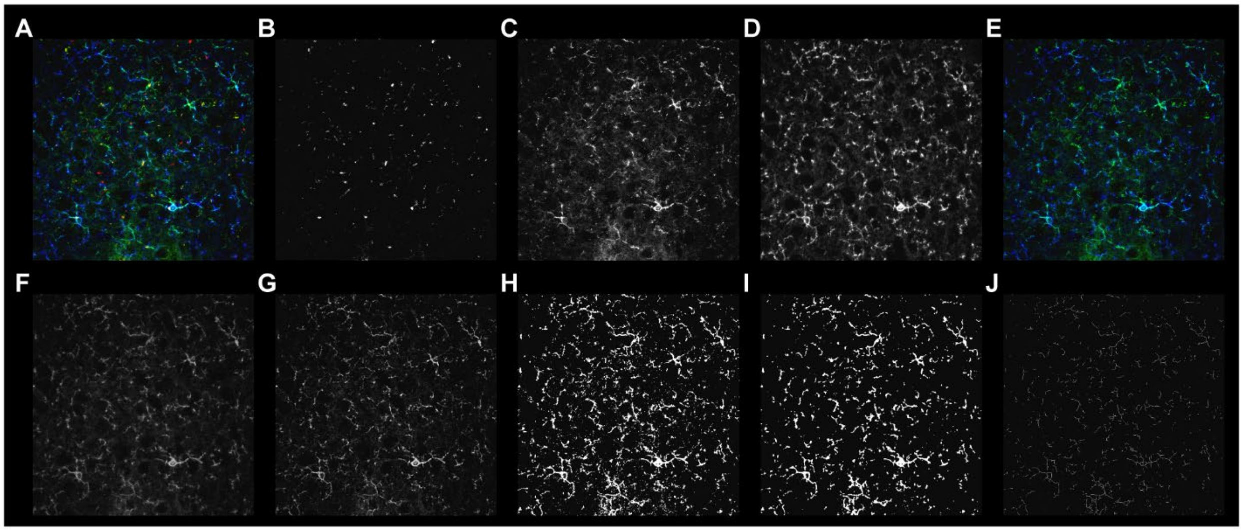

**Fig. S1: Representative micrographs for Morphology image analysis pipeline.** For printing purposes, all images are shown as Max Intensity Z-Projections. **A**. Original multi-channel image. **B**. Red channel from original image (A). **C**. Green channel from original image (A). **D**. Blue channel from original image (A). **E**. Green (C) and blue (D) channels merged. **F**. Merged image (E) converted to grayscale. **G**. Grayscale image (F) after application of Unsharp Mask and Despeckle tools. **H**. De-noised image (G) after thresholding. **I**. Thresholded image (H) after application of Despeckle, Close-, and Remove Outliers tools. **J**. Final skeletonized image used to quantify branching.

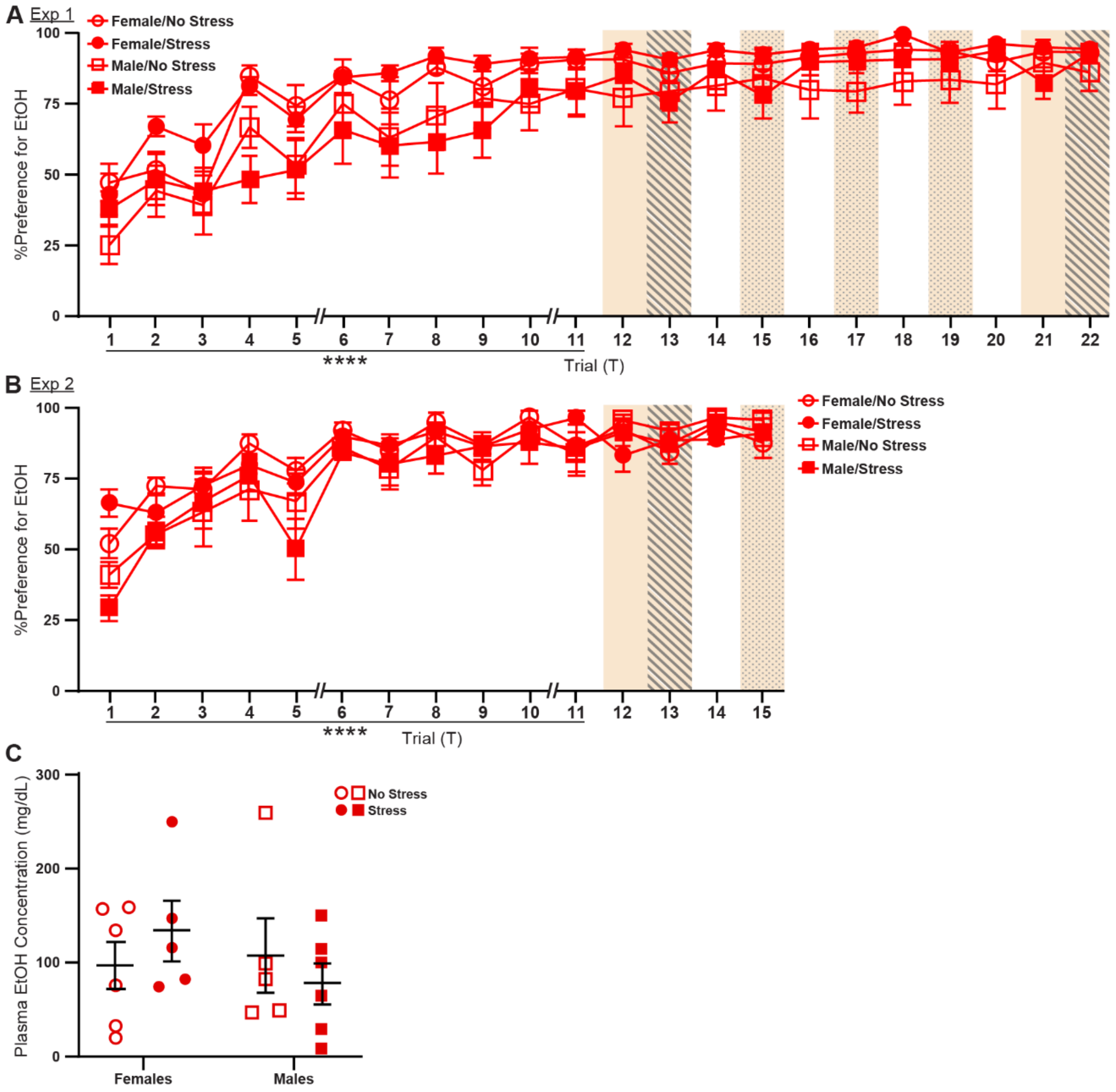

**Fig. S2: Additional characterization of behavioral changes in Experiments 1 and 2, related to Fig. 1.**

**A**. Time course of relative preference for ethanol per subgroup across 22 trials of DiD for Experiment 1. All mice increased ethanol preference over the course of the acquisition period (trials 1-11). There were no significant differences in relative preference for ethanol after trial 11. **B**. Mice in Experiment 2 showed similar trends of relative preference for ethanol across the acquisition period. **C**. Plasma EtOH concentrations after the last 4 hr drinking trial in a separate cohort of mice that followed the same design as Experiment 1. There were no statistical differences among experimental groups. **p* < 0.0001; *n* = 11/group. Data represent average +/- SEM.

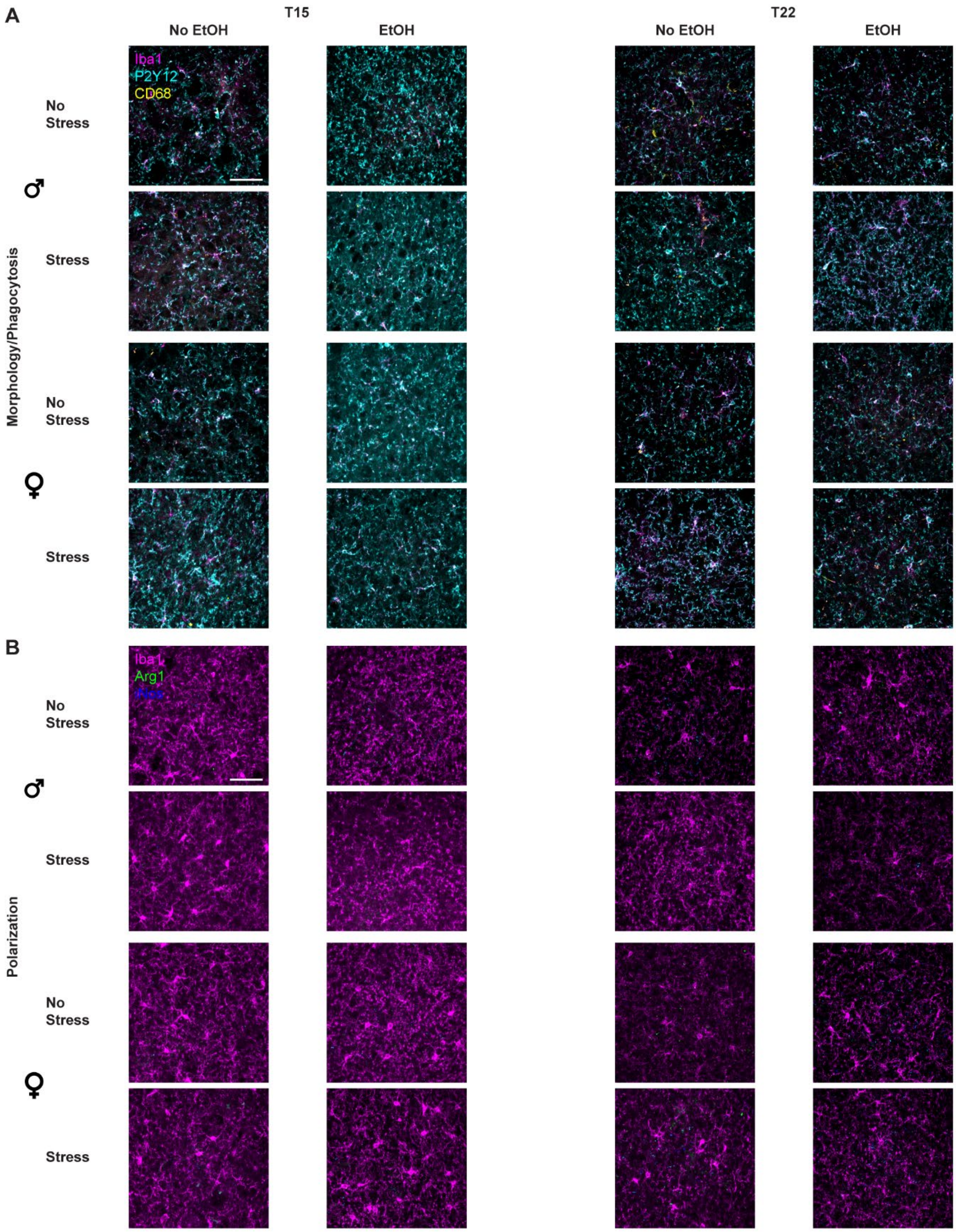

**Fig. S3: Representative micrographs for BLA microglia in Experiments 1 & 2, related to Fig. 2.**

All images are shown as Max Intensity Z-Projections. **A**. Representative images for the Morphology/Phagocytosis stain; Iba1 is magenta, CD68 is yellow, and P2Y12 is cyan.

**B**. Representative images for the Polarization stain; Iba1 is magenta, Arg1 is green, and iNos is blue. Scale bar = 50 µm.

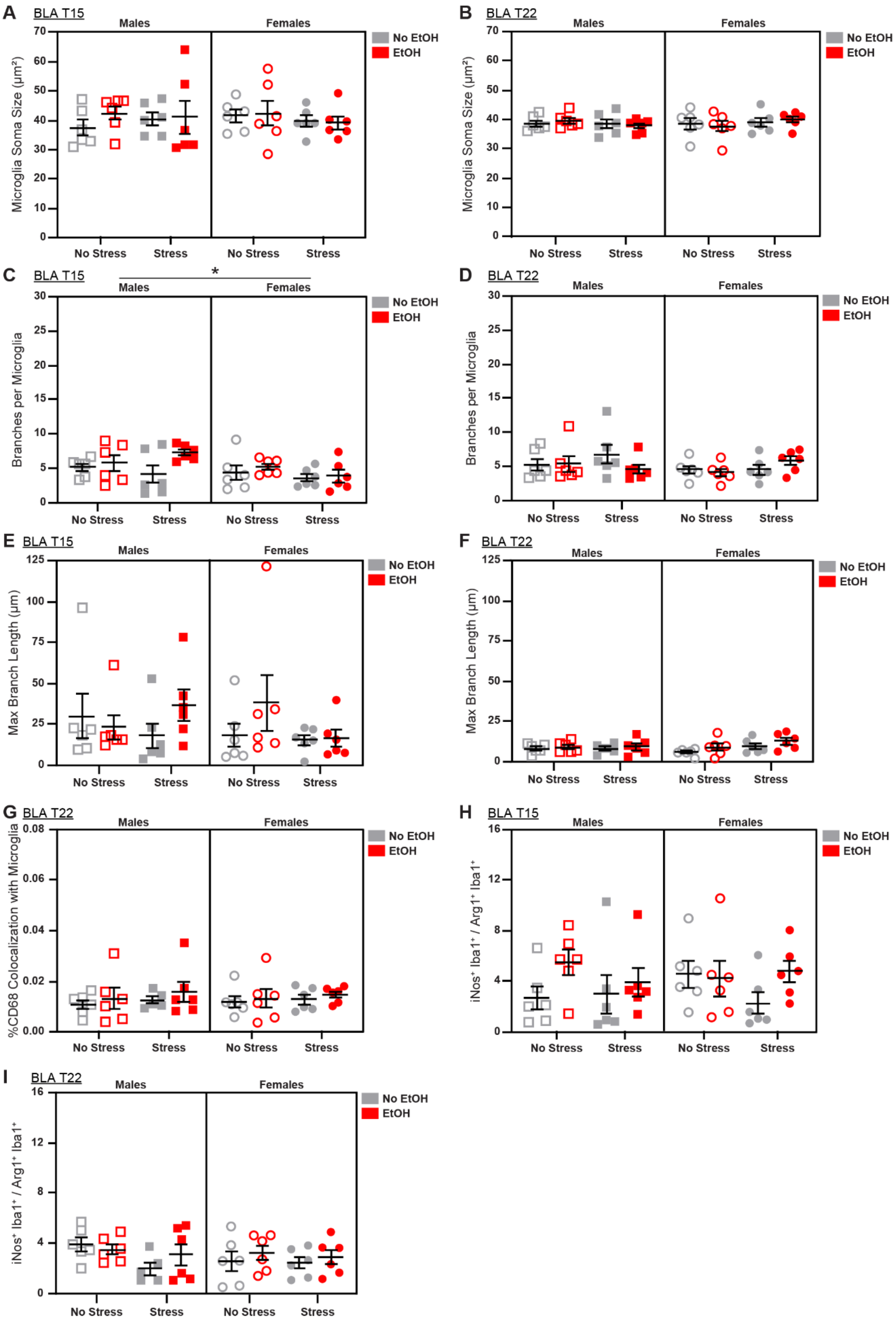
A
BLA T15
Males
Females
No EtOH
EtOH
Microglia Soma Size (μm²)
No Stress
Stress
B
BLA T22
C
BLA T15
*
Branches per Microglia
D
BLA T22
E
BLA T15
Max Branch Length (μm)
F
BLA T22
G
BLA T22
%CD68 Colocalization with Microglia
H
BLA T15
iNos⁺ Iba1⁺ / Arg1⁺ Iba1⁺
I
BLA T22

**Fig. S4: Additional characterization of sex, stress, and EtOH effects on microglial phenotypes in BLA from Experiments 1 & 2, related to Fig. 2**
**A**. There were no main effects of sex, stress, or EtOH on microglia soma size on T15, **B.** or T22. **C.** Branch number was increased in males on T15, **D.** but this effect did not persist to T22. **E.** There were no main effects of sex, stress, or EtOH on branch length on T15, **F.** or T22, **G.** or on microglial CD68 expression on T22 (1 outlier excluded). **H.** There were no main effects of sex, stress, or EtOH on M1/M2 polarization on T15, **I.** or T22 (1 outlier excluded). See Fig. S3 for representative micrographs and Table S2 for detailed statistics. *$p < 0.05$; $n$ = 5-6 animals/group.

**Table S2: Complete statistical analyses for characterization of BLA microglia in Experiments 1 & 2, related to Fig. 2.**
Data shown in Figs. 2 and S4. *$p < 0.05$.

| | **T15** | | **T22** | |
|---|---|---|---|---|
| **Analysis** | ***F / t*** | ***p*** | ***F / t*** | ***p*** |
| 3-way ANOVA: Microglia Density | | | | |
| Effect of Sex | $F_{1,40}$ = 2.827 | 0.1005 | $F_{1,40}$ = 0.1979 | 0.6588 |
| Effect of Stress | $F_{1,40}$ = 2.246 | 0.1418 | $F_{1,40}$ = 0.7380 | 0.3954 |
| Effect of EtOH | $F_{1,40}$ = 5.776 | 0.0210* | $F_{1,40}$ = 0.3213 | 0.5740 |
| Sex x Stress Interaction | $F_{1,40}$ = 1.448 | 0.2359 | $F_{1,40}$ = 0.8070 | 0.3744 |
| Sex x EtOH Interaction | $F_{1,40}$ = 0.09070 | 0.7648 | $F_{1,40}$ = 5.425 | 0.0250* |
| Stress x EtOH Interaction | $F_{1,40}$ = 0.1668 | 0.6851 | $F_{1,40}$ = 0.05048 | 0.8234 |
| Sex x Stress x EtOH Interaction | $F_{1,40}$ = 6.278 | 0.0164* | $F_{1,40}$ = 2.990 | 0.0915 |
| 2-way ANOVA: Microglia Density, Males | | | | |
| Effect of Stress | $F_{1,20}$ = 0.04283 | 0.8381 | $F_{1,20}$ = 0.001002 | 0.2396 |
| Effect of EtOH | $F_{1,20}$ = 3.598 | 0.0724 | $F_{1,20}$ = 5.443 | 0.0302* |
| Stress x EtOH Interaction | $F_{1,20}$ = 4.178 | 0.0544 | $F_{1,20}$ = 1.469 | 0.2396 |
| Šídák's multiple comparisons test: Microglia Density, Males | | | | |
| No Stress/No EtOH v. No Stress/EtOH | | | $t$ = 0.7927 | 0.6833 |
| Stress/No EtOH v. Stress/EtOH | | | $t$ = 2.507 | 0.0414* |
| No EtOH/No Stress v. No EtOH/Stress | | | $t$ = 0.8346 | 0.6563 |

| | | | | |
|---|---|---|---|---|
| EtOH/No Stress v. EtOH/Stress | | | $t$ = 0.8794 | 0.6274 |
| 2-way ANOVA: Microglia Density, Females | | | | |
| Effect of Stress | $F_{1,20}$ = 3.711 | 0.0684 | $F_{1,20}$ = 1.256 | 0.2757 |
| Effect of EtOH | $F_{1,20}$ = 2.246 | 0.1496 | $F_{1,20}$ = 1.263 | 0.2744 |
| Stress x EtOH Interaction | $F_{1,20}$ = 2.236 | 0.1505 | $F_{1,20}$ = 1.552 | 0.2272 |
| 3-way ANOVA: Microglia Soma Size | | | | |
| Effect of Sex | $F_{1,40}$ = 0.0228 | 0.8807 | $F_{1,40}$ = 0.01323 | 0.9090 |
| Effect of Stress | $F_{1,40}$ = 0.1240 | 0.7266 | $F_{1,40}$ = 0.02411 | 0.8774 |
| Effect of EtOH | $F_{1,40}$ = 0.4541 | 0.5043 | $F_{1,40}$ = 0.01529 | 0.9022 |
| Sex x Stress Interaction | $F_{1,40}$ = 0.5633 | 0.4573 | $F_{1,40}$ = 1.308 | 0.2595 |
| Sex x EtOH Interaction | $F_{1,40}$ = 0.3285 | 0.5697 | $F_{1,40}$ = 0.01099 | 0.9170 |
| Stress x EtOH Interaction | $F_{1,40}$ = 0.4078 | 0.5267 | $F_{1,40}$ = 0.001190 | 0.9727 |
| Sex x Stress x EtOH Interaction | $F_{1,40}$ = 0.09806 | 0.7558 | $F_{1,40}$ = 0.8471 | 0.3629 |
| 3-way ANOVA: Branches per Microglia | | | | |
| Effect of Sex | $F_{1,40}$ = 4.683 | 0.0365* | $F_{1,40}$ = 1.601 | 0.2131 |
| Effect of Stress | $F_{1,40}$ = 0.4065 | 0.5274 | $F_{1,40}$ = 0.9688 | 0.3309 |
| Effect of EtOH | $F_{1,40}$ = 4.021 | 0.0517 | $F_{1,40}$ = 0.2696 | 0.6065 |
| Sex x Stress Interaction | $F_{1,40}$ = 1.079 | 0.3052 | $F_{1,40}$ = 0.1434 | 0.7069 |
| Sex x EtOH Interaction | $F_{1,40}$ = 1.117 | 0.2970 | $F_{1,40}$ = 1.513 | 0.2259 |
| Stress x EtOH Interaction | $F_{1,40}$ = 0.5978 | 0.440 | $F_{1,40}$ = 0.06967 | 0.7932 |
| Sex x Stress x EtOH Interaction | $F_{1,40}$ = 1.785 | 0.1891 | $F_{1,40}$ = 2.521 | 0.1202 |
| 2-way ANOVA: Branches per Microglia, Males | | | | |
| Effect of Stress | $F_{1,20}$ = 0.06825 | 0.7966 | | |
| Effect of EtOH | $F_{1,20}$ = 3.979 | 0.0599 | | |
| Stress x EtOH Interaction | $F_{1,20}$ = 1.888 | 0.1846 | | |

| | | | | |
|---|---|---|---|---|
| 2-way ANOVA: Branches per Microglia, Females | | | | |
| Effect of Stress | $F_{1,20}$ = 1.709 | 0.2059 | | |
| Effect of EtOH | $F_{1,20}$ = 0.5473 | 0.4680 | | |
| Stress x EtOH Interaction | $F_{1,20}$ = 0.1927 | 0.6654 | | |
| 3-way ANOVA: Max Branch Length | | | | |
| Effect of Sex | $F_{1,40}$ = 0.5019 | 0.4828 | $F_{1,39}$ = 0.2489 | 0.6206 |
| Effect of Stress | $F_{1,40}$ = 0.6967 | 0.4088 | $F_{1,39}$ = 2.733 | 0.1063 |
| Effect of EtOH | $F_{1,40}$ = 1.415 | 0.2413 | $F_{1,39}$ = 3.269 | 0.0783 |
| Sex x Stress Interaction | $F_{1,40}$ = 0.8789 | 0.3541 | $F_{1,39}$ = 2.330 | 0.1350 |
| Sex x EtOH Interaction | $F_{1,40}$ = 0.09262 | 0.7625 | $F_{1,39}$ = 0.8869 | 0.3521 |
| Stress x EtOH Interaction | $F_{1,40}$ = 0.05124 | 0.8221 | $F_{1,39}$ = 0.0003573 | 0.9850 |
| Sex x Stress x EtOH Interaction | $F_{1,40}$ = 2.526 | 0.1198 | $F_{1,39}$ = 0.004653 | 0.9460 |
| 3-way ANOVA: %CD68 Colocalization with Microglia | | | | |
| Effect of Sex | $F_{1,40}$ = 0.09250 | 0.7626 | $F_{1,39}$ = 0.0008815 | 0.9765 |
| Effect of Stress | $F_{1,40}$ = 0.03130 | 0.8605 | $F_{1,39}$ = 0.7014 | 0.4074 |
| Effect of EtOH | $F_{1,40}$ = 6.936 | 0.0120* | $F_{1,39}$ = 1.137 | 2.928 |
| Sex x Stress Interaction | $F_{1,40}$ = 0.1541 | 0.6967 | $F_{1,39}$ = 0.05490 | 0.8160 |
| Sex x EtOH Interaction | $F_{1,40}$ = 0.1403 | 0.7099 | $F_{1,39}$ = 0.08755 | 0.7689 |
| Stress x EtOH Interaction | $F_{1,40}$ = 0.004071 | 0.9494 | $F_{1,39}$ = 0.04733 | 0.8289 |
| Sex x Stress x EtOH Interaction | $F_{1,40}$ = 0.02871 | 0.8663 | $F_{1,39}$ = 0.002687 | 0.9589 |
| 2-way ANOVA: %CD68 Colocalization with Microglia, Males | | | | |
| Effect of Stress | $F_{1,20}$ = 0.02358 | 0.8795 | | |
| Effect of EtOH | $F_{1,20}$ = 4.588 | 0.0447* | | |
| Stress x EtOH Interaction | $F_{1,20}$ = 0.02759 | 0.8697 | | |

| | | | | |
|---|---|---|---|---|
| Šídák's multiple comparisons test: %CD68 Colocalization with Microglia, Males | | | | |
| No Stress/No EtOH v. No Stress/EtOH | $t$ = 1.397 | 0.3238 | | |
| Stress/No EtOH v. Stress/EtOH | $t$ = 1.632 | 0.2226 | | |
| No EtOH/No Stress v. No EtOH/Stress | $t$ = 0.2260 | 0.9688 | | |
| EtOH/No Stress v. EtOH/Stress | $t$ = 0.008869 | >0.9999 | | |
| 2-way ANOVA: %CD68 Colocalization with Microglia, Females | | | | |
| Effect of Stress | $F_{1,20}$ = 0.1599 | 0.6935 | | |
| Effect of EtOH | $F_{1,20}$ = 2.516 | 0.1284 | | |
| Stress x EtOH Interaction | $F_{1,20}$ = 0.005504 | 0.9416 | | |
| 3-way ANOVA: iNos$^+$ Iba1$^+$ / Arg1$^+$ Iba1$^+$ | | | | |
| Effect of Sex | $F_{1,40}$ = 0.06263 | 0.8037 | $F_{1,39}$ = 0.6208 | 0.4355 |
| Effect of Stress | $F_{1,40}$ = 0.9220 | 0.3427 | $F_{1,39}$ = 2.710 | 0.1077 |
| Effect of EtOH | $F_{1,40}$ = 3.656 | 0.0630 | $F_{1,39}$ = 1.135 | 0.2933 |
| Sex x Stress Interaction | $F_{1,40}$ = 0.02014 | 0.8879 | $F_{1,39}$ = 1.146 | 0.2910 |
| Sex x EtOH Interaction | $F_{1,40}$ = 0.2647 | 0.6097 | $F_{1,39}$ = 0.04913 | 0.8257 |
| Stress x EtOH Interaction | $F_{1,40}$ = 0.08034 | 0.7783 | $F_{1,39}$ = 0.6369 | 0.4297 |
| Sex x Stress x EtOH Interaction | $F_{1,40}$ = 2.309 | 0.1365 | $F_{1,39}$ = 1.017 | 0.3194 |

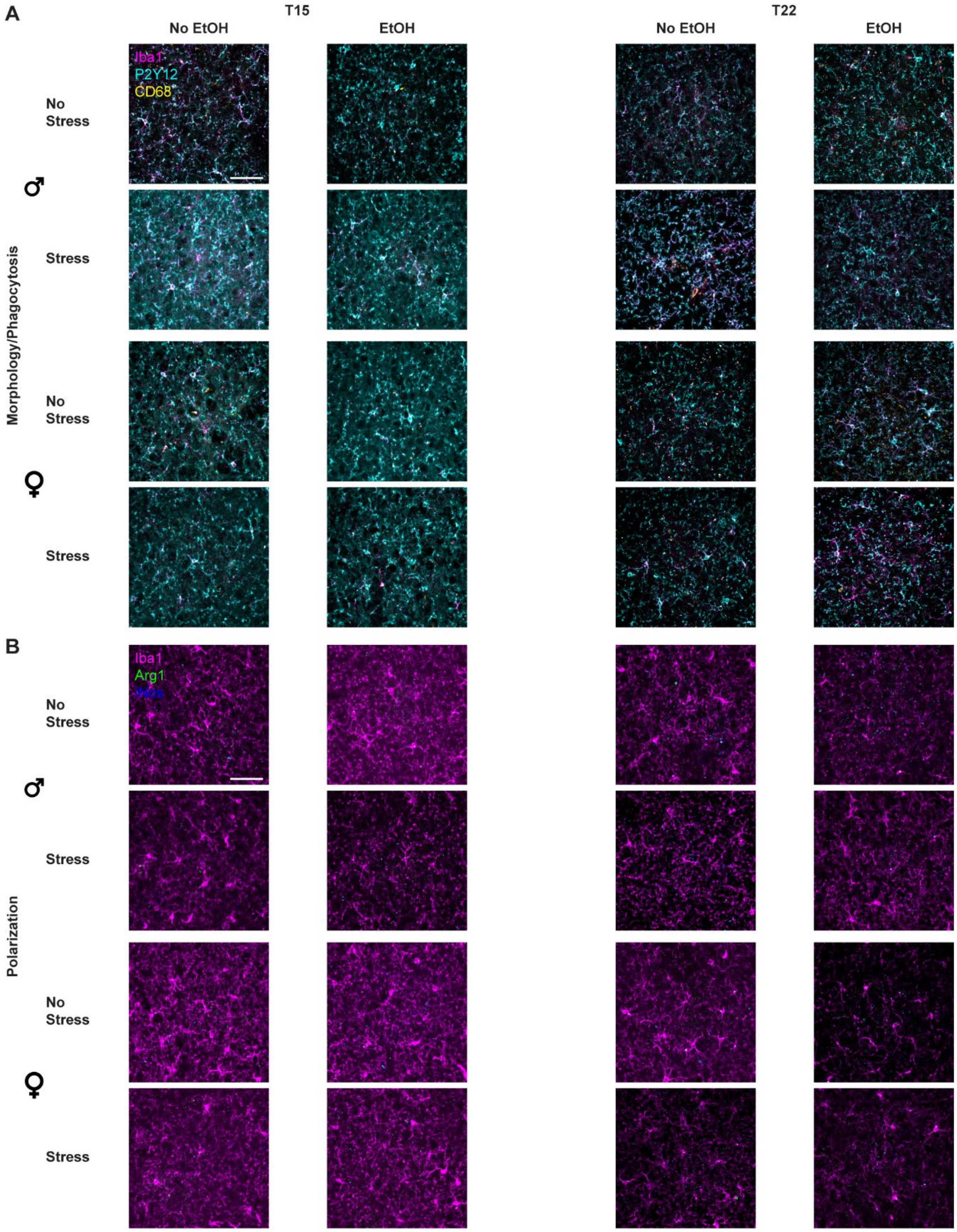

**Fig. S5: Representative micrographs for CeA microglia in Experiments 1 & 2, related to Fig. 2.**

All images are shown as Max Intensity Z-Projections. **A**. Representative images for the Morphology/Phagocytosis stain; Iba1 is magenta, CD68 is yellow, and P2Y12 is cyan.

**B**. Representative images for the Polarization stain; Iba1 is magenta, Arg1 is green, and iNos is blue. Scale bar = 50 μm.

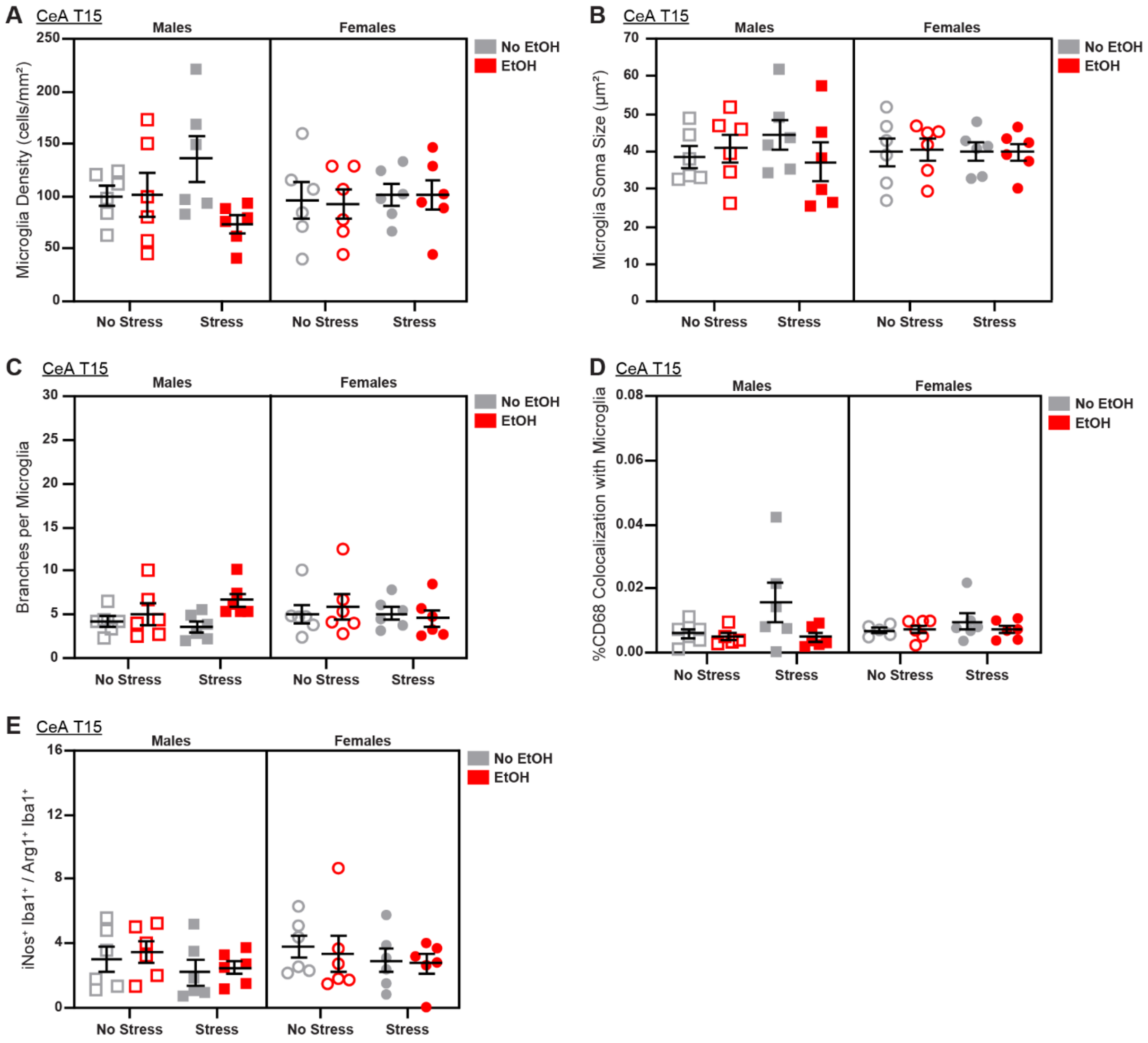

**Fig. S6: Additional characterization of sex, stress, and EtOH effects on microglial phenotypes in CeA from Experiment 2, related to Fig. 2**
At T15, there were no main effects of sex, stress, or EtOH on **A**. Microglia density, **B.** microglia soma size, **C.** branch length, **D.** microglial CD68 expression (2 outliers excluded), **E.** or M1/M2 polarization. See Fig. S5 for representative micrographs and Table S3 for detailed statistics. *n* = 5-6 animals/group.

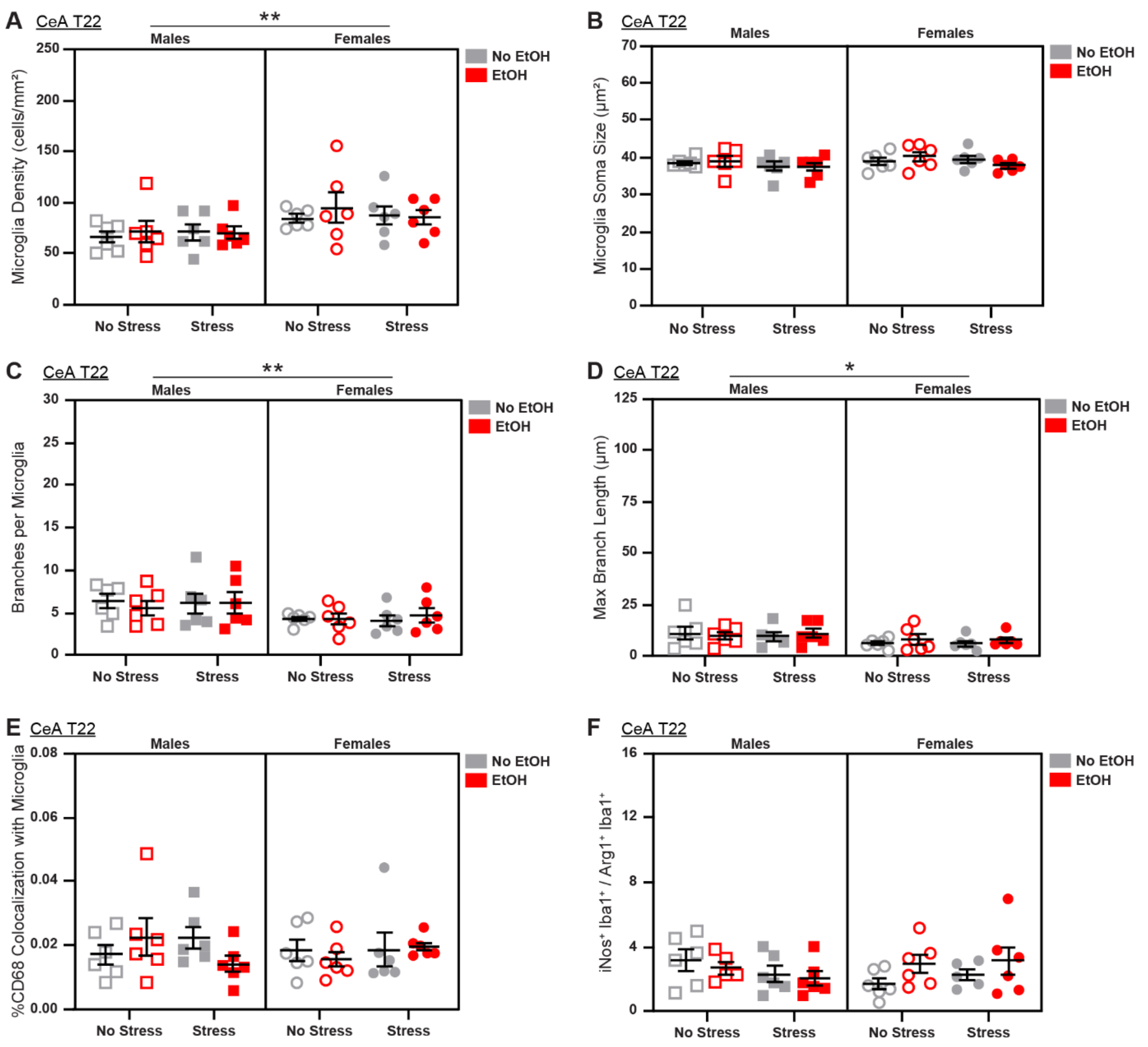

**Fig. S7: Additional characterization of sex, stress, and EtOH effects on microglial phenotypes in CeA from Experiment 1, related to Fig. 2**

At T22, **A.** microglial density was increased in females. **B.** There were no main effects of sex, stress, or EtOH on microglia soma size (1 outlier excluded). **C.** Males showed increased branch number **D.** and branch length (1 outlier excluded). **E.** There were no main effects of sex, stress, or EtOH on microglial CD68 expression or **F.** M1/M2 polarization (2 outliers excluded). See Fig. S5 for representative micrographs and Table S3 for detailed statistics. *$p$ < 0.05; **$p$ < 0.01; $n$ = 5-6 animals/group.

**Table S3: Complete statistical analyses for characterization of CeA microglia in Experiments 1 & 2, related to Fig. 2**
Data shown in Figs. 2 and S6-7. *$p < 0.05$; **$p < 0.01$.

| | T15 | | T22 | |
|---|---|---|---|---|
| **Analysis** | ***F / t*** | ***p*** | ***F / t*** | ***p*** |
| 3-way ANOVA: Microglia Density | | | | |
| Effect of Sex | $F_{1,40}$ = 0.2003 | 0.6569 | $F_{1,40}$ = 8.986 | 0.0047** |
| Effect of Stress | $F_{1,40}$ = 0.2487 | 0.6208 | $F_{1,40}$ = 0.01877 | 0.8917 |
| Effect of EtOH | $F_{1,40}$ = 2.315 | 0.1360 | $F_{1,40}$ = 0.2865 | 0.5955 |
| Sex x Stress Interaction | $F_{1,40}$ = 0.01746 | 0.8955 | $F_{1,40}$ = 0.1664 | 0.6855 |
| Sex x EtOH Interaction | $F_{1,40}$ = 1.734 | 0.1954 | $F_{1,40}$ = 0.01974 | 0.8890 |
| Stress x EtOH Interaction | $F_{1,40}$ = 1.919 | 0.1737 | $F_{1,40}$ = 0.6193 | 0.4359 |
| Sex x Stress x EtOH Interaction | $F_{1,40}$ = 2.420 | 0.1277 | $F_{1,40}$ = 0.06705 | 0.7970 |
| 2-way ANOVA: Microglia Density, Males | | | | |
| Effect of Stress | | | $F_{1,20}$ = 0.04924 | 0.8266 |
| Effect of EtOH | | | $F_{1,20}$ = 0.1046 | 0.7498 |
| Stress x EtOH Interaction | | | $F_{1,20}$ =0.1871 | 0.6700 |
| 2-way ANOVA: Microglia Density, Females | | | | |
| Effect of Stress | | | $F_{1,20}$ = 0.1183 | 0.7345 |
| Effect of EtOH | | | $F_{1,20}$ = 0.1819 | 0.6743 |
| Stress x EtOH Interaction | | | $F_{1,20}$ = 0.4358 | 0.5167 |
| 3-way ANOVA: Microglia Soma Size | | | | |
| Effect of Sex | $F_{1,40}$ = 0.007658 | 0.9307 | $F_{1,40}$ = 1.818 | 0.1852 |
| Effect of Stress | $F_{1,40}$ = 0.02466 | 0.8760 | $F_{1,40}$ = 2.250 | 0.1415 |
| Effect of EtOH | $F_{1,40}$ = 0.1873 | 0.6675 | $F_{1,40}$ = 0.002722 | 0.9587 |

| | | | | |
|---|---|---|---|---|
| Sex x Stress Interaction | $F_{1,40}$ = 0.08601 | 0.7708 | $F_{1,40}$ = 0.06918 | 0.7939 |
| Sex x EtOH Interaction | $F_{1,40}$ = 0.3238 | 0.5725 | $F_{1,40}$ = 0.09593 | 0.7584 |
| Stress x EtOH Interaction | $F_{1,40}$ = 1.063 | 0.3087 | $F_{1,40}$ = 1.435 | 0.2380 |
| Sex x Stress x EtOH Interaction | $F_{1,40}$ = 0.7935 | 0.3784 | $F_{1,40}$ = 0.9271 | 0.3414 |
| 3-way ANOVA: Branches per Microglia | | | | |
| Effect of Sex | $F_{1,40}$ = 0.2268 | 0.6365 | $F_{1,40}$ = 7.910 | 0.0076** |
| Effect of Stress | $F_{1,40}$ = 0.01084 | 0.9176 | $F_{1,40}$ = 0.05415 | 0.8172 |
| Effect of EtOH | $F_{1,40}$ = 2.481 | 0.1231 | $F_{1,40}$ = 0.001023 | 0.9746 |
| Sex x Stress Interaction | $F_{1,40}$ = 0.7438 | 0.3936 | $F_{1,40}$ = 0.0002639 | 0.9871 |
| Sex x EtOH Interaction | $F_{1,40}$ = 1.885 | 0.1774 | $F_{1,40}$ = 0.2593 | 0.6134 |
| Stress x EtOH Interaction | $F_{1,40}$ = 0.09391 | 0.7608 | $F_{1,40}$ = 0.3519 | 0.5564 |
| Sex x Stress x EtOH Interaction | $F_{1,40}$ = 1.713 | 0.1981 | $F_{1,40}$ = 0.00417 | 0.9473 |
| 2-way ANOVA: Branches per Microglia, Males | | | | |
| Effect of Stress | | | $F_{1,20}$ = 0.02124 | 0.8856 |
| Effect of EtOH | | | $F_{1,20}$ = 0.07808 | 0.7828 |
| Stress x EtOH Interaction | | | $F_{1,20}$ = 0.1492 | 0.7034 |
| 2-way ANOVA: Branches per Microglia, Females | | | | |
| Effect of Stress | | | $F_{1,20}$ = 0.04326 | 0.8373 |
| Effect of EtOH | | | $F_{1,20}$ = 0.2705 | 0.6087 |
| Stress x EtOH Interaction | | | $F_{1,20}$ = 0.2562 | 0.6183 |
| 3-way ANOVA: Max Branch Length | | | | |
| Effect of Sex | $F_{1,40}$ = 0.004799 | 0.9451 | $F_{1,39}$ = 0.5371 | 0.0258* |

| | | | | |
|---|---|---|---|---|
| Effect of Stress | $F_{1,40}$ = 0.001569 | 0.9686 | $F_{1,39}$ = 0.02632 | 0.8720 |
| Effect of EtOH | $F_{1,40}$ = 4.727 | 0.0357* | $F_{1,39}$ = 0.2620 | 0.6117 |
| Sex x Stress Interaction | $F_{1,40}$ = 2.479 | 0.1232 | $F_{1,39}$ = 0.002680 | 0.9590 |
| Sex x EtOH Interaction | $F_{1,40}$ = 1.351 | 0.2520 | $F_{1,39}$ = 0.4144 | 0.5235 |
| Stress x EtOH Interaction | $F_{1,40}$ = 1.231 | 0.2738 | $F_{1,39}$ = 0.1276 | 0.7229 |
| Sex x Stress x EtOH Interaction | $F_{1,40}$ = 0.5198 | 0.4751 | $F_{1,39}$ = 0.2882 | 0.5944 |
| 2-way ANOVA: Max Branch Length, Males | | | | |
| Effect of Stress | $F_{1,20}$ = 1.185 | 0.2893 | $F_{1,19}$ = 0.004239 | 0.9488 |
| Effect of EtOH | $F_{1,20}$ = 5.063 | 0.0359* | $F_{1,19}$ = 0.006046 | 0.9388 |
| Stress x EtOH Interaction | $F_{1,20}$ = 1.524 | 0.2313 | $F_{1,19}$ = 0.2777 | 0.6043 |
| Šídák’s multiple comparisons test: %CD68 Colocalization with Microglia, Males | | | | |
| No Stress/No EtOH v. No Stress/EtOH | *t* = 0.7181 | 0.7306 | | |
| Stress/No EtOH v. Stress/EtOH | *t* = 2.464 | 0.0453* | | |
| No EtOH/No Stress v. No EtOH/Stress | *t* = 0.1032 | 0.9934 | | |
| EtOH/No Stress v. EtOH/Stress | *t* = 1.643 | 0.2187 | | |
| 2-way ANOVA: Max Branch Length, Females | | | | |
| Effect of Stress | $F_{1,20}$ = 1.308 | 0.2663 | $F_{1,20}$ = 0.03815 | 0.8471 |
| Effect of EtOH | $F_{1,20}$ = 0.5687 | 0.4596 | $F_{1,20}$ = 1.112 | 0.3042 |
| Stress x EtOH Interaction | $F_{1,20}$ = 0.08386 | 0.7751 | $F_{1,20}$ = 0.02687 | 0.8714 |
| 3-way ANOVA: %CD68 Colocalization with Microglia | | | | |
| Effect of Sex | $F_{1,38}$ = 0.001451 | 0.9698 | $F_{1,40}$ = 0.1337 | 0.7166 |
| Effect of Stress | $F_{1,38}$ = 2.592 | 0.1157 | $F_{1,40}$ = 0.007200 | 0.9328 |

| | | | | |
|---|---|---|---|---|
| Effect of EtOH | $F_{1,38}$ = 3.463 | 0.0705 | $F_{1,40}$ = 0.2066 | 0.6519 |
| Sex x Stress Interaction | $F_{1,38}$ = 0.8684 | 0.3573 | $F_{1,40}$ = 0.5560 | 0.4602 |
| Sex x EtOH Interaction | $F_{1,38}$ = 1.673 | 0.2036 | $F_{1,40}$ = 0.004394 | 0.9475 |
| Stress x EtOH Interaction | $F_{1,38}$ = 2.935 | 0.0948 | $F_{1,40}$ = 0.9009 | 0.3482 |
| Sex x Stress x EtOH Interaction | $F_{1,38}$ = 0.8739 | 0.3558 | $F_{1,40}$ = 2.936 | 0.0944 |
| 3-way ANOVA: $iNos^+$ $Iba1^+$ / $Arg1^+$ $Iba1^+$ | | | | |
| Effect of Sex | $F_{1,40}$ = 0.5891 | 0.4473 | $F_{1,38}$ = 0.02071 | 0.8863 |
| Effect of Stress | $F_{1,40}$ = 2.518 | 0.1204 | $F_{1,38}$ = 0.2375 | 0.6288 |
| Effect of EtOH | $F_{1,40}$ = 0.004818 | 0.9450 | $F_{1,38}$ = 0.7223 | 0.4007 |
| Sex x Stress Interaction | $F_{1,40}$ = 0.02018 | 0.8878 | $F_{1,38}$ = 2.187 | 0.1474 |
| Sex x EtOH Interaction | $F_{1,40}$ = 0.4629 | 0.5002 | $F_{1,38}$ = 3.405 | 0.0728 |
| Stress x EtOH Interaction | $F_{1,40}$ = 0.001456 | 0.9698 | $F_{1,38}$ = 0.013818 | 0.9092 |
| Sex x Stress x EtOH Interaction | $F_{1,40}$ = 0.04654 | 0.8303 | $F_{1,38}$ = 0.1227 | 0.7281 |

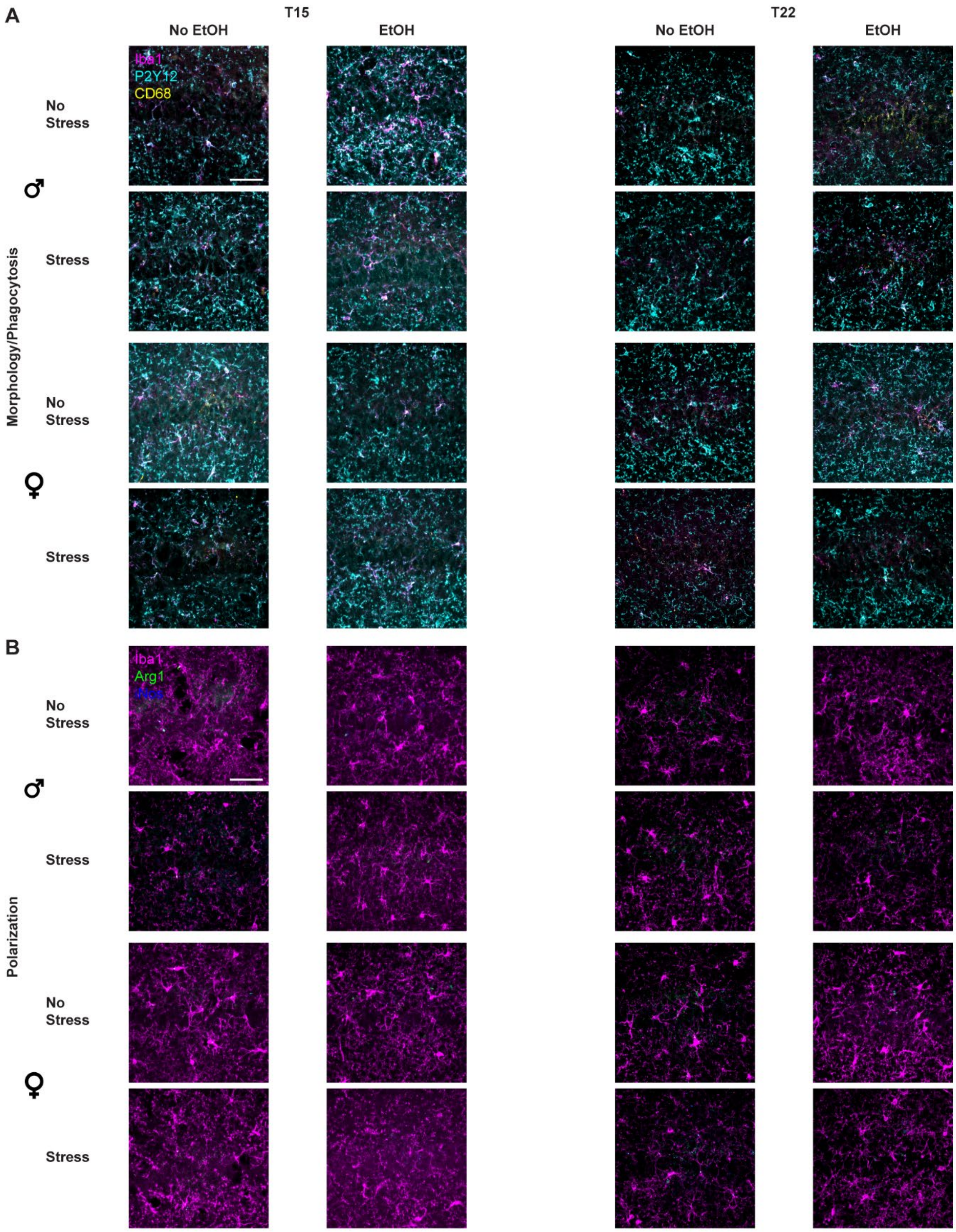

**Fig. S8: Representative micrographs for CA1 microglia in Experiments 1 & 2, related to Fig. 3.**

All images are shown as Max Intensity Z-Projections. **A**. Representative images for the Morphology/Phagocytosis stain; Iba1 is magenta, CD68 is yellow, and P2Y12 is cyan.

**B**. Representative images for the Polarization stain; Iba1 is magenta, Arg1 is green, and iNos is blue. Scale bar = 50 μm.

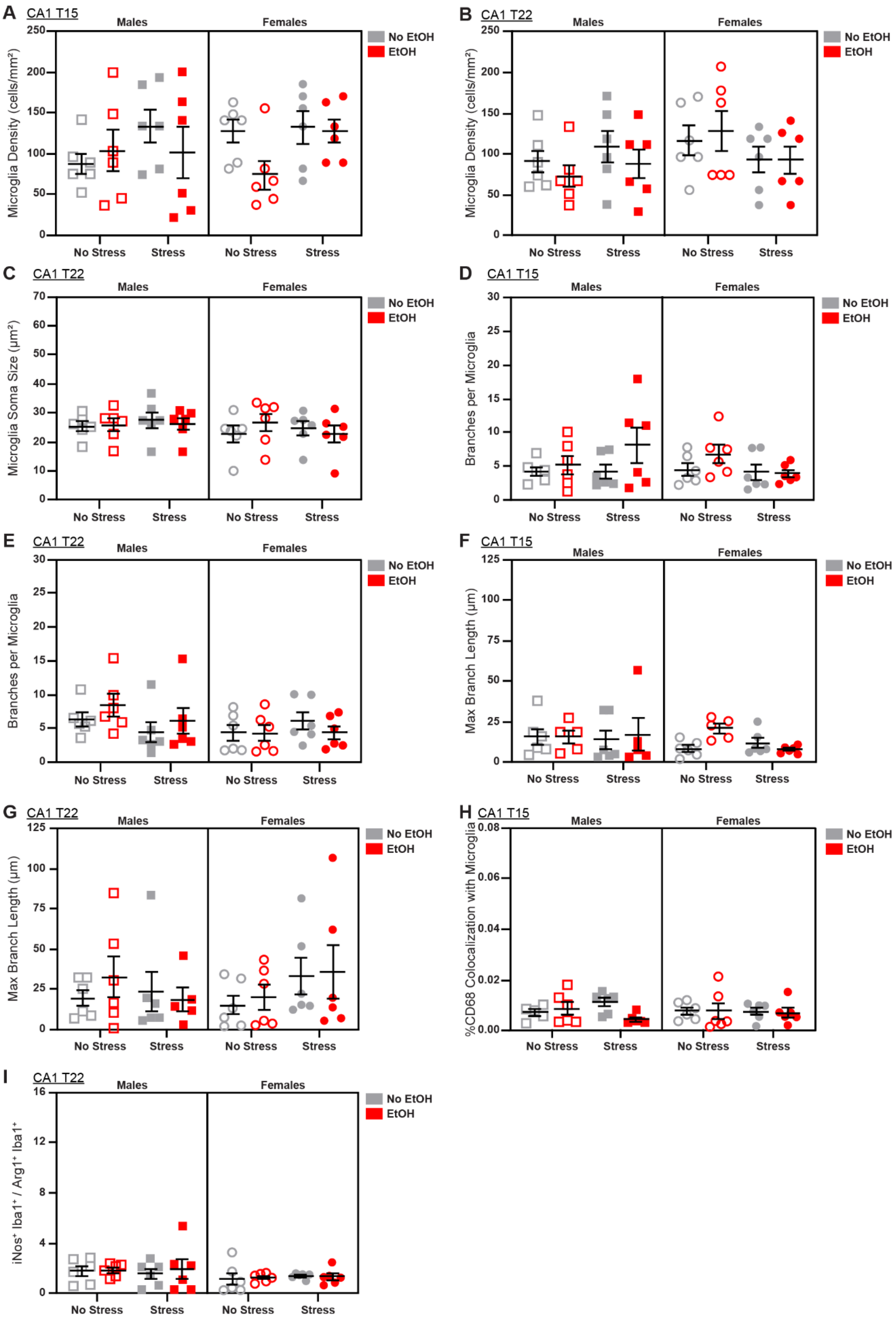
A
CA1 T15
B
CA1 T22
C
CA1 T22
D
CA1 T15
E
CA1 T22
F
CA1 T15
G
CA1 T22
H
CA1 T15
I
CA1 T22
Males
Females
No EtOH
EtOH
No Stress
Stress
Microglia Density (cells/mm²)
Microglia Soma Size (µm²)
Branches per Microglia
Max Branch Length (µm)
%CD68 Colocalization with Microglia
iNos⁺ Iba1⁺ / Arg1⁺ Iba1⁺

**Fig. S9: Additional characterization of sex, stress, and EtOH effects on microglial phenotypes in CA1 from Experiments 1 & 2, related to Fig. 3**
There were no main effects of sex, stress, or EtOH on **A**. microglia density on T15, **B.** microglia density on T22, **C.** microglia soma size on T22, **D.** branch number on T15 (1 outlier excluded), **E.** branch number on T22, **F.** branch length on T15 (4 outliers excluded), **G.** branch length on T22 (1 outlier excluded), **H.** microglial CD68 expression on T15 (2 outliers excluded), or **I.** M1/M2 polarization on T22 (1 outlier excluded). See Fig. S8 for representative micrographs and Table S4 for detailed statistics. *n* = 5-6 animals/group.

**Table S4: Complete statistical analyses for characterization of CA1 microglia in Experiments 1 & 2, related to Fig. 3.**
Data shown in Figs. 3 and S9. $*p < 0.05$.

| | T15 | | T22 | |
|---|---|---|---|---|
| **Analysis** | ***F / t*** | ***p*** | ***F / t*** | ***p*** |
| 3-way ANOVA: Microglia Density | | | | |
| Effect of Sex | $F_{1,40}$ = 0.3647 | 0.5488 | $F_{1,40}$ = 1.977 | 0.1674 |
| Effect of Stress | $F_{1,40}$ = 3.136 | 0.0842 | $F_{1,40}$ = 0.2721 | 0.6048 |
| Effect of EtOH | $F_{1,40}$ = 1.679 | 0.2025 | $F_{1,40}$ = 0.2982 | 0.5881 |
| Sex x Stress Interaction | $F_{1,40}$ = 0.06717 | 0.7968 | $F_{1,40}$ = 3.282 | 0.0776 |
| Sex x EtOH Interaction | $F_{1,40}$ = 0.5392 | 0.4671 | $F_{1,40}$ = 0.9961 | 0.3243 |
| Stress x EtOH Interaction | $F_{1,40}$ = 1.846e-014 | >0.9999 | $F_{1,40}$ = 0.1017 | 0.7515 |
| Sex x Stress x EtOH Interaction | $F_{1,40}$ = 2.838 | 0.0999 | $F_{1,40}$ = 0.03313 | 0.8565 |
| 3-way ANOVA: Microglia Soma Size | | | | |
| Effect of Sex | $F_{1,40}$ = 0.8704 | 0.3564 | $F_{1,40}$ = 1.291 | 0.2625 |
| Effect of Stress | $F_{1,40}$ = 0.6902 | 0.4110 | $F_{1,40}$ = 0.008168 | 0.9284 |
| Effect of EtOH | $F_{1,40}$ = 0.5732 | 0.4534 | $F_{1,40}$ = 0.02822 | 0.8674 |
| Sex x Stress Interaction | $F_{1,40}$ = 5.739 | 0.0214* | $F_{1,40}$ = 0.3882 | 0.5368 |
| Sex x EtOH Interaction | $F_{1,40}$ = 0.007586 | 0.9310 | $F_{1,40}$ = 0.1634 | 0.6882 |
| Stress x EtOH Interaction | $F_{1,40}$ = 0.6302 | 0.4320 | $F_{1,40}$ = 1.135 | 0.2931 |
| Sex x Stress x EtOH Interaction | $F_{1,40}$ = 0.01932 | 0.8901 | $F_{1,40}$ = 0.3199 | 0.5748 |

| | | | | |
|---|---|---|---|---|
| 2-way ANOVA: Microglia Soma Size, Males | | | | |
| Effect of Stress | $F_{1,20}$ = 4.543 | 0.0456* | | |
| Effect of EtOH | $F_{1,20}$ = 0.3110 | 0.5832 | | |
| Stress x EtOH Interaction | $F_{1,20}$ = 0.3797 | 0.5447 | | |
| Šídák's multiple comparisons test: Microglia Soma Size, Males | | | | |
| No Stress/No EtOH v. No Stress/EtOH | $t$ = 0.04139 | 0.9989 | | |
| Stress/No EtOH v. Stress/EtOH | $t$ = 0.8301 | 0.6593 | | |
| No EtOH/No Stress v. No EtOH/Stress | $t$ = 1.071 | 0.5054 | | |
| EtOH/No Stress v. EtOH/Stress | $t$ = 1.943 | 0.1281 | | |
| 2-way ANOVA: Microglia Soma Size, Females | | | | |
| Effect of Stress | $F_{1,20}$ = 1.433 | 0.2452 | | |
| Effect of EtOH | $F_{1,20}$ = 0.2628 | 0.6138 | | |
| Stress x EtOH Interaction | $F_{1,20}$ = 0.2510 | 0.6219 | | |
| 3-way ANOVA: Branches per Microglia | | | | |
| Effect of Sex | $F_{1,40}$ = 0.4230 | 0.5192 | $F_{1,40}$ = 2.512 | 0.1209 |
| Effect of Stress | $F_{1,40}$ = 0.005900 | 0.9392 | $F_{1,40}$ = 0.3860 | 0.5379 |
| Effect of EtOH | $F_{1,40}$ = 3.431 | 0.0714 | $F_{1,40}$ = 0.2549 | 0.6164 |
| Sex x Stress Interaction | $F_{1,40}$ = 2.682 | 0.1093 | $F_{1,40}$ = 2.460 | 0.1247 |
| Sex x EtOH Interaction | $F_{1,40}$ = 0.6160 | 0.4372 | $F_{1,40}$ = 1.959 | 0.1693 |
| Stress x EtOH Interaction | $F_{1,40}$ = 0.007748 | 0.9303 | $F_{1,40}$ = 0.3119 | 0.5796 |
| Sex x Stress x EtOH Interaction | $F_{1,40}$ = 2.154 | 0.1500 | $F_{1,40}$ = 0.09718 | 0.7569 |
| 3-way ANOVA: Max Branch Length | | | | |
| Effect of Sex | $F_{1,37}$ = 1.051 | 0.3119 | $F_{1,39}$ = 0.1128 | 0.7388 |
| Effect of Stress | $F_{1,37}$ = 0.5785 | 0.4517 | $F_{1,39}$ = 0.6162 | 0.4372 |
| Effect of EtOH | $F_{1,37}$ = 0.7525 | 0.3913 | $F_{1,39}$ = 0.2892 | 0.5938 |
| Sex x Stress Interaction | $F_{1,37}$ = 0.4718 | 0.4964 | $F_{1,39}$ = 2.057 | 0.1595 |

| | | | | |
|---|---|---|---|---|
| Sex x EtOH Interaction | $F_{1,37}$ = 0.1723 | 0.6805 | $F_{1,39}$ = 0.0007983 | 0.9776 |
| Stress x EtOH Interaction | $F_{1,37}$ = 1.048 | 0.3126 | $F_{1,39}$ = 0.4326 | 0.5146 |
| Sex x Stress x EtOH Interaction | $F_{1,37}$ = 2.055 | 0.1601 | $F_{1,39}$ = 0.2749 | 0.6030 |
| 3-way ANOVA: %CD68 Colocalization with Microglia | | | | |
| Effect of Sex | $F_{1,39}$ = 0.06064 | 0.8068 | $F_{1,38}$ = 0.3313 | 0.5683 |
| Effect of Stress | $F_{1,39}$ = 0.04270 | 0.8374 | $F_{1,38}$ = 0.7376 | 0.3958 |
| Effect of EtOH | $F_{1,39}$ = 1.056 | 0.3104 | $F_{1,38}$ = 0.7009 | 0.4077 |
| Sex x Stress Interaction | $F_{1,39}$ = 0.01309 | 0.9095 | $F_{1,38}$ = 0.1877 | 0.6673 |
| Sex x EtOH Interaction | $F_{1,39}$ = 0.7188 | 0.4017 | $F_{1,38}$ = 7.100 | 0.0113* |
| Stress x EtOH Interaction | $F_{1,39}$ = 2.509 | 0.1213 | $F_{1,38}$ = 0.1301 | 0.7203 |
| Sex x Stress x EtOH Interaction | $F_{1,39}$ = 2.023 | 0.1629 | $F_{1,38}$ = 3.455 | 0.0708 |
| 2-way ANOVA: %CD68 Colocalization with Microglia, Males | | | | |
| Effect of Stress | | | $F_{1,19}$ = 0.1030 | 0.7518 |
| Effect of EtOH | | | $F_{1,19}$ = 6.971 | 0.0161* |
| Stress x EtOH Interaction | | | $F_{1,19}$ = 1.276 | 0.2728 |
| Šídák's multiple comparisons test: %CD68 Colocalization with Microglia, Males | | | | |
| No Stress/No EtOH v. No Stress/EtOH | | | $t$ = 1.095 | 0.4921 |
| Stress/No EtOH v. Stress/EtOH | | | $t$ = 2.604 | 0.0345* |
| No EtOH/No Stress v. No EtOH/Stress | | | $t$ = 1.002 | 0.5497 |
| EtOH/No Stress v. EtOH/Stress | | | $t$ = 0.5858 | 0.8107 |
| 2-way ANOVA: %CD68 Colocalization with Microglia, Females | | | | |
| Effect of Stress | | | $F_{1,19}$ = 0.7449 | 0.3989 |

| | | | | |
|---|---|---|---|---|
| Effect of EtOH | | | $F_{1,19}$ = 1.490 | 0.2371 |
| Stress x EtOH Interaction | | | $F_{1,19}$ = 2.198 | 0.1546 |
| 3-way ANOVA: iNos$^+$ Iba1$^+$ / Arg1$^+$ Iba1$^+$ | | | | |
| Effect of Sex | $F_{1,40}$ = 0.1981 | 0.6586 | $F_{1,40}$ = 3.073 | 0.0873 |
| Effect of Stress | $F_{1,40}$ = 0.6327 | 0.4311 | $F_{1,40}$ = 0.02317 | 0.8798 |
| Effect of EtOH | $F_{1,40}$ = 1.697 | 0.2002 | $F_{1,40}$ = 0.1769 | 0.6763 |
| Sex x Stress Interaction | $F_{1,40}$ = 0.1915 | 0.6640 | $F_{1,40}$ = 0.1379 | 0.7123 |
| Sex x EtOH Interaction | $F_{1,40}$ = 0.1235 | 0.7272 | $F_{1,40}$ = 0.1220 | 0.7288 |
| Stress x EtOH Interaction | $F_{1,40}$ = 0.04656 | 0.8303 | $F_{1,40}$ = 0.01493 | 0.9033 |
| Sex x Stress x EtOH Interaction | $F_{1,40}$ = 4.120 | 0.0491* | $F_{1,40}$ = 0.1993 | 0.6577 |
| 2-way ANOVA: iNos$^+$ Iba1$^+$ / Arg1$^+$ Iba1$^+$, Males | | | | |
| Effect of Stress | $F_{1,20}$ = 0.04373 | 0.8365 | | |
| Effect of EtOH | $F_{1,20}$ = 0.9346 | 0.3452 | | |
| Stress x EtOH Interaction | $F_{1,20}$ = 1.723 | 0.2042 | | |
| 2-way ANOVA: iNos$^+$ Iba1$^+$ / Arg1$^+$ Iba1$^+$, Females | | | | |
| Effect of Stress | $F_{1,20}$ = 1.417 | 0.2479 | | |
| Effect of EtOH | $F_{1,20}$ = 0.8431 | 0.3695 | | |
| Stress x EtOH Interaction | $F_{1,20}$ = 3.066 | 0.0953 | | |

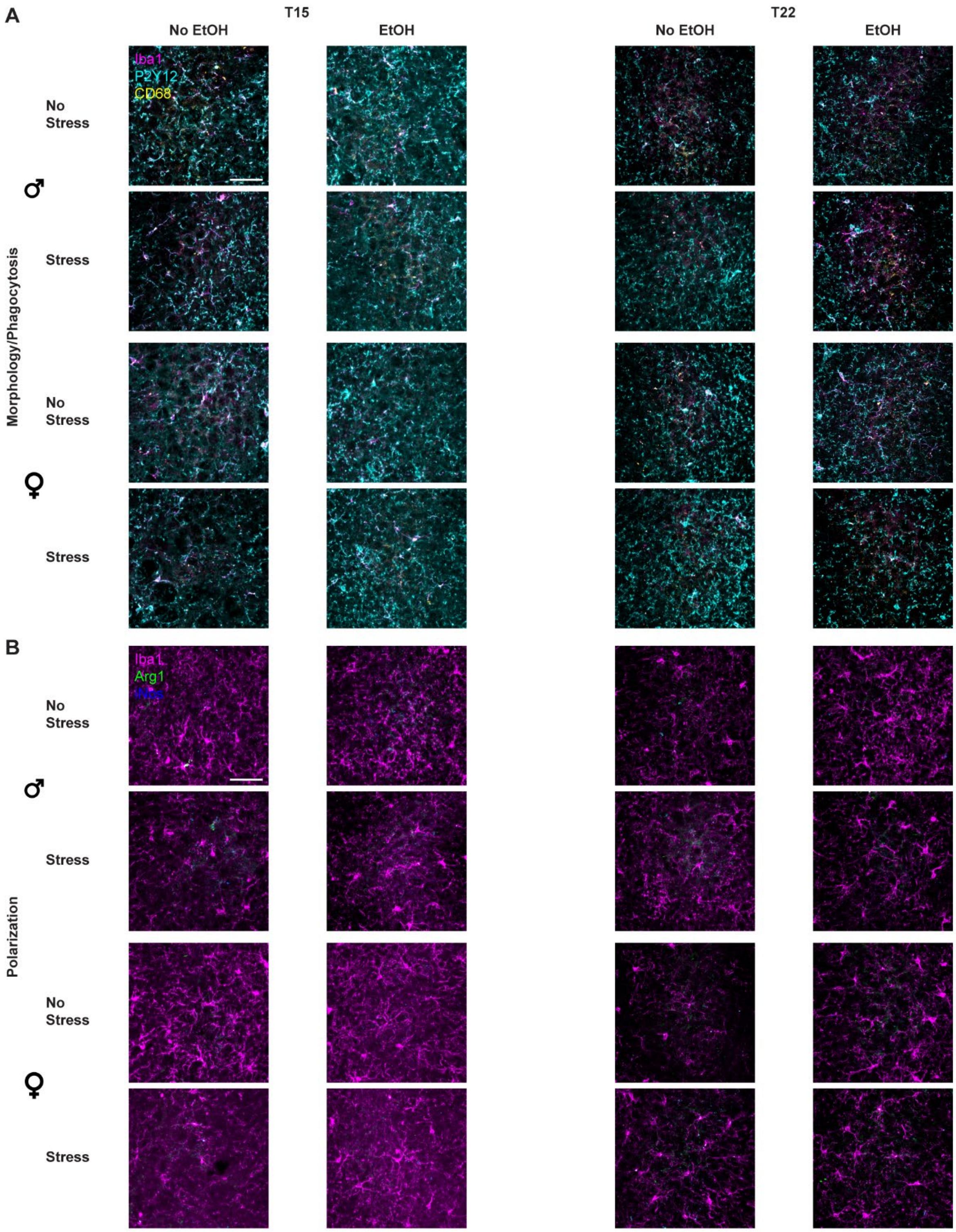

**Fig. S10: Representative micrographs for CA3 microglia in Experiments 1 & 2, related to Fig. 3.**

All images are shown as Max Intensity Z-Projections. **A**. Representative images for the Morphology/Phagocytosis stain; Iba1 is magenta, CD68 is yellow, and P2Y12 is cyan.

**B**. Representative images for the Polarization stain; Iba1 is magenta, Arg1 is green, and iNos is blue. Scale bar = 50 μm.

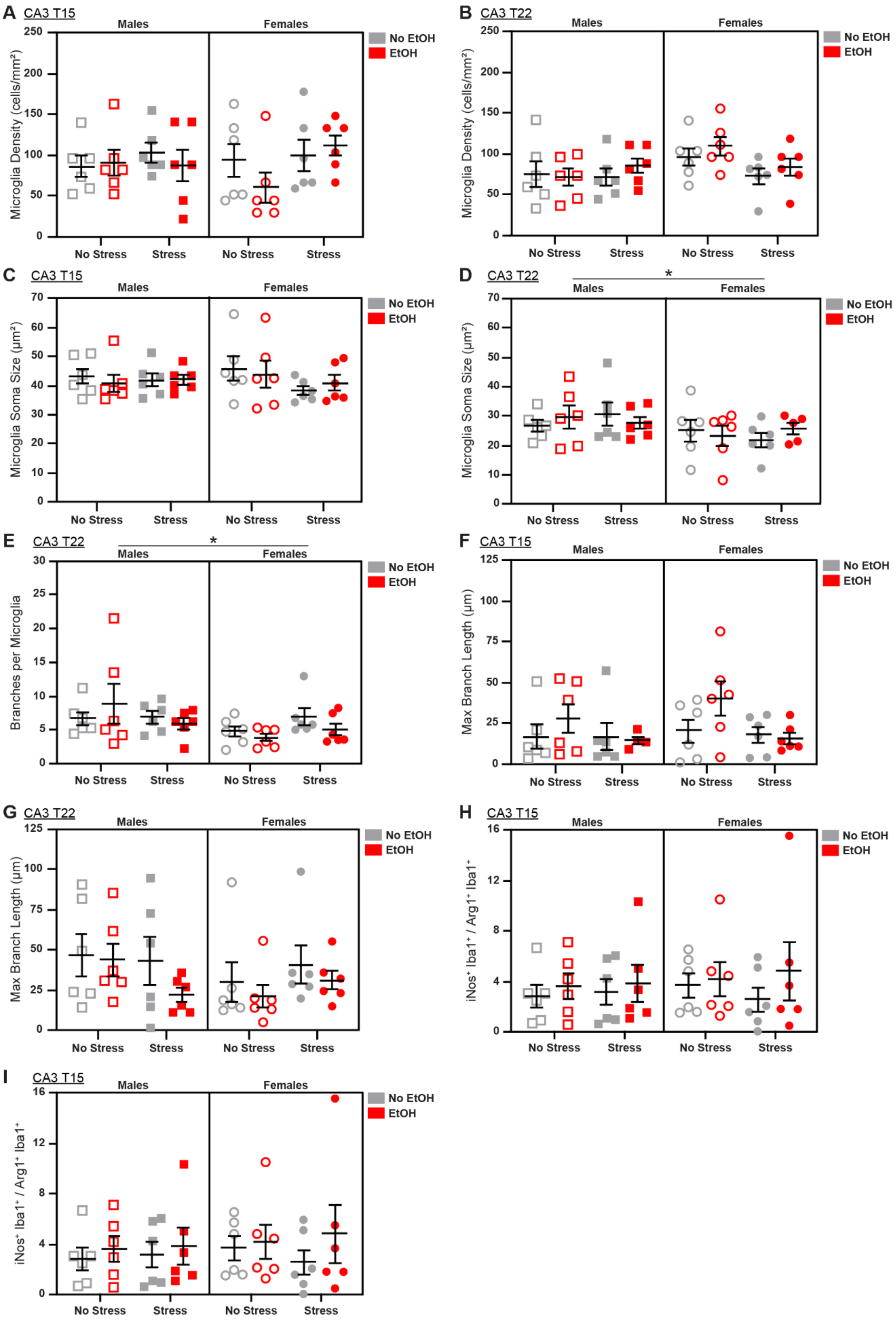

A
CA3 T15
Males
Females
No EtOH
EtOH
Microglia Density (cells/mm²)
250
200
150
100
50
0
No Stress
Stress
B
CA3 T22
Microglia Density (cells/mm²)
C
CA3 T15
Microglia Soma Size (µm²)
70
60
40
30
20
10
D
CA3 T22
*
Microglia Soma Size (µm²)
E
CA3 T22
*
Branches per Microglia
25
15
5
F
CA3 T15
Max Branch Length (µm)
125
75
25
G
CA3 T22
Max Branch Length (µm)
H
CA3 T15
iNos⁺ Iba1⁺ / Arg1⁺ Iba1⁺
16
12
8
4
I
CA3 T15
iNos⁺ Iba1⁺ / Arg1⁺ Iba1⁺

**Fig. S11: Additional characterization of sex, stress, and EtOH effects on microglial phenotypes in CA3 from Experiments 1 & 2, related to Fig. 3**
There were no main effects of sex, stress, or EtOH on **A**. microglia density on T15, **B.** microglia density on T22, **C.** or microglia soma size on T15. **D.** On T22, males show increased soma size (1 outlier excluded), **E.** and branch number. **F.** There were no main effects of sex, stress, or EtOH on branch length at T15 (2 outliers excluded), **G.** branch length on T22, **H.** M1/M2 polarization on T15, **I.** or M1/M2 polarization on T22. See Fig. S10 for representative micrographs and Table S5 for detailed statistics. *$p < 0.05$; $n$ = 4-6 animals/group.

**Table S5: Complete statistical analyses for characterization of CA3 microglia in Experiments 1 & 2, related to Fig. 3.**
Data shown in Figs. 3 and S11. *$p < 0.05$; **$p < 0.01$.

| | T15 | | T22 | |
|---|---|---|---|---|
| **Analysis** | ***F / t*** | ***p*** | ***F / t*** | ***p*** |
| 3-way ANOVA: Microglia Density | | | | |
| Effect of Sex | $F_{1,40}$ = 0.0006745 | 0.9794 | $F_{1,40}$ = 3.389 | 0.0730 |
| Effect of Stress | $F_{1,40}$ = 2.348 | 0.1333 | $F_{1,40}$ = 1.406 | 0.2428 |
| Effect of EtOH | $F_{1,40}$ = 0.4917 | 0.4872 | $F_{1,40}$ = 1.186 | 0.2826 |
| Sex x Stress Interaction | $F_{1,40}$ = 0.8262 | 0.3688 | $F_{1,40}$ = 3.461 | 0.0702 |
| Sex x EtOH Interaction | $F_{1,40}$ = 0.03305 | 0.8567 | $F_{1,40}$ = 0.2416 | 0.6257 |
| Stress x EtOH Interaction | $F_{1,40}$ = 0.2975 | 0.5885 | $F_{1,40}$ = 0.2610 | 0.6123 |
| Sex x Stress x EtOH Interaction | $F_{1,40}$ = 1.895 | 0.1763 | $F_{1,40}$ = 0.4423 | 0.5098 |
| 3-way ANOVA: Microglia Soma Size | | | | |
| Effect of Sex | $F_{1,40}$ = 0.006672 | 0.9353 | $F_{1,39}$ = 4.507 | 0.0401* |
| Effect of Stress | $F_{1,40}$ = 1.661 | 0.2049 | $F_{1,39}$ = 0.01782 | 0.8945 |
| Effect of EtOH | $F_{1,40}$ = 0.02800 | 0.8680 | $F_{1,39}$ = 0.06973 | 0.7931 |
| Sex x Stress Interaction | $F_{1,40}$ = 1.482 | 0.2306 | $F_{1,39}$ = 0.1079 | 0.7443 |
| Sex x EtOH Interaction | $F_{1,40}$ = 0.1182 | 0.7328 | $F_{1,39}$ = 0.04457 | 0.8339 |
| Stress x EtOH Interaction | $F_{1,40}$ = 0.6810 | 0.4141 | $F_{1,39}$ = 0.0004148 | 0.9839 |

| | | | | |
|---|---|---|---|---|
| Sex x Stress x EtOH Interaction | $F_{1,40}$ = 0.06622 | 0.7982 | $F_{1,39}$ = 1.647 | 0.2069 |
| 2-way ANOVA: Microglia Soma Size, Males | | | | |
| Effect of Stress | | | $F_{1,20}$ = 1.063 | 0.7478 |
| Effect of EtOH | | | $F_{1,20}$ = 0.001396 | 0.9706 |
| Stress x EtOH Interaction | | | $F_{1,20}$ = 0.8467 | 0.3684 |
| 2-way ANOVA: Microglia Soma Size, Females | | | | |
| Effect of Stress | | | $F_{1,19}$ = 0.01913 | 0.8915 |
| Effect of EtOH | | | $F_{1,19}$ = 0.1136 | 0.7397 |
| Stress x EtOH Interaction | | | $F_{1,19}$ = 0.8029 | 0.3814 |
| 3-way ANOVA: Branches per Microglia | | | | |
| Effect of Sex | $F_{1,39}$ = 2.134 | 0.1521 | $F_{1,40}$ = 4.178 | 0.0476* |
| Effect of Stress | $F_{1,39}$ = 4.647 | 0.0373* | $F_{1,40}$ = 0.01562 | 0.9012 |
| Effect of EtOH | $F_{1,39}$ = 0.1377 | 0.7126 | $F_{1,40}$ = 0.1537 | 0.6971 |
| Sex x Stress Interaction | $F_{1,39}$ = 3.972 | 0.0533 | $F_{1,40}$ = 2.637 | 0.1122 |
| Sex x EtOH Interaction | $F_{1,39}$ = 0.01717 | 0.8964 | $F_{1,40}$ = 1.127 | 0.2949 |
| Stress x EtOH Interaction | $F_{1,39}$ = 0.2004 | 0.6569 | $F_{1,40}$ = 1.219 | 0.2761 |
| Sex x Stress x EtOH Interaction | $F_{1,39}$ = 2.720 | 0.1071 | $F_{1,40}$ = 0.3580 | 0.5530 |
| 2-way ANOVA: Branches per Microglia, Males | | | | |
| Effect of Stress | $F_{1,19}$ = 0.01807 | 0.8945 | $F_{1,20}$ = 0.7283 | 0.4035 |
| Effect of EtOH | $F_{1,19}$ = 0.03923 | 0.8451 | $F_{1,20}$ = 0.1452 | 0.7072 |
| Stress x EtOH Interaction | $F_{1,19}$ = 0.9833 | 0.3339 | $F_{1,20}$ = 0.9394 | 0.3440 |
| 2-way ANOVA: Branches per Microglia, Females | | | | |
| Effect of Stress | $F_{1,20}$ = 6.950 | 0.0158* | $F_{1,20}$ = 0.2797 | 0.6027 |
| Effect of EtOH | $F_{1,20}$ = 0.1018 | 0.7530 | $F_{1,20}$ = 3.345 | 0.0823 |

| | | | | |
|---|---|---|---|---|
| Stress x EtOH Interaction | $F_{1,20}$ = 1.776 | 0.1977 | $F_{1,20}$ = 2.310 | 0.1442 |
| Šídák's multiple comparisons test: Branches per Microglia, Females | | | | |
| No Stress/No EtOH v. No Stress/EtOH | $t$ = 1.168 | 0.4474 | | |
| Stress/No EtOH v. Stress/EtOH | $t$ = 0.7166 | 0.7316 | | |
| No EtOH/No Stress v. No EtOH/Stress | $t$ = 0.9219 | 0.6000 | | |
| EtOH/No Stress v. EtOH/Stress | $t$ = 2.806 | 0.0217* | | |
| 3-way ANOVA: Max Branch Length | | | | |
| Effect of Sex | $F_{1,40}$ = 0.7343 | 0.3969 | $F_{1,40}$ = 1.192 | 0.2814 |
| Effect of Stress | $F_{1,40}$ = 3.731 | 0.0609 | $F_{1,40}$ = 0.02974 | 0.8639 |
| Effect of EtOH | $F_{1,40}$ = 1.655 | 0.2061 | $F_{1,40}$ = 2.067 | 0.1583 |
| Sex x Stress Interaction | $F_{1,40}$ = 0.4035 | 0.5291 | $F_{1,40}$ = 2.425 | 0.1273 |
| Sex x EtOH Interaction | $F_{1,40}$ = 0.1710 | 0.6815 | $F_{1,40}$ = 0.03629 | 0.8499 |
| Stress x EtOH Interaction | $F_{1,40}$ = 2.869 | 0.0985 | $F_{1,40}$ = 0.4011 | 0.5301 |
| Sex x Stress x EtOH Interaction | $F_{1,40}$ = 0.1448 | 0.7056 | $F_{1,40}$ = 0.3214 | 0.5739 |
| 3-way ANOVA: %CD68 Colocalization with Microglia | | | | |
| Effect of Sex | $F_{1,40}$ = 5.442 | 0.0248* | $F_{1,39}$ = 9.9685 | 0.3311 |
| Effect of Stress | $F_{1,40}$ = 0.04512 | 0.8329 | $F_{1,39}$ = 6.434 | 0.0153* |
| Effect of EtOH | $F_{1,40}$ = 7.581 | 0.0088** | $F_{1,39}$ = 0.4048 | 0.5283 |
| Sex x Stress Interaction | $F_{1,40}$ = 0.003077 | 0.9560 | $F_{1,39}$ = 0.8974 | 0.3493 |
| Sex x EtOH Interaction | $F_{1,40}$ = 0.04127 | 0.8400 | $F_{1,39}$ = 1.529 | 0.2237 |
| Stress x EtOH Interaction | $F_{1,40}$ = 0.3798 | 0.5412 | $F_{1,39}$ = 1.824e-005 | 0.9966 |
| Sex x Stress x EtOH Interaction | $F_{1,40}$ = 0.1705 | 0.6818 | $F_{1,39}$ = 1.257 | 0.2691 |

| | | | | |
|---|---|---|---|---|
| 2-way ANOVA: %CD68 Colocalization with Microglia, Males | | | | |
| Effect of Stress | $F_{1,20}$ = 0.02517 | 0.8755 | $F_{1,20}$ = 3.113 | 0.0929 |
| Effect of EtOH | $F_{1,20}$ = 3.066 | 0.0953 | $F_{1,20}$ = 0.4438 | 0.5129 |
| Stress x EtOH Interaction | $F_{1,20}$ = 0.3716 | 0.5490 | $F_{1,20}$ = 1.537 | 0.2294 |
| 2-way ANOVA: %CD68 Colocalization with Microglia, Females | | | | |
| Effect of Stress | $F_{1,20}$ = 0.02144 | 0.8850 | $F_{1,19}$ = 3.669 | 0.0706 |
| Effect of EtOH | $F_{1,20}$ = 5.661 | 0.0274* | $F_{1,19}$ = 1.060 | 0.3162 |
| Stress x EtOH Interaction | $F_{1,20}$ = 0.03599 | 0.8514 | $F_{1,19}$ = 0.3828 | 0.5435 |
| Šídák’s multiple comparisons test: %CD68 Colocalization with Microglia, Females | | | | |
| No Stress/No EtOH v. No Stress/EtOH | *t* = 1.817 | 0.1615 | | |
| Stress/No EtOH v. Stress/EtOH | *t* = 1.548 | 0.2556 | | |
| No EtOH/No Stress v. No EtOH/Stress | *t* = 0.2377 | 0.9656 | | |
| EtOH/No Stress v. EtOH/Stress | *t* = 0.03061 | 0.9994 | | |
| 3-way ANOVA: $iNos^{+}$ $Iba1^{+}$ / $Arg1^{+}$ $Iba1^{+}$ | | | | |
| Effect of Sex | $F_{1,40}$ = 0.2493 | 0.6203 | $F_{1,40}$ = 0.4670 | 0.4983 |
| Effect of Stress | $F_{1,40}$ = 0.0001753 | 0.9895 | $F_{1,40}$ = 2.003 | 0.1648 |
| Effect of EtOH | $F_{1,40}$ = 1.319 | 0.2576 | $F_{1,40}$ = 0.004019 | 0.9498 |
| Sex x Stress Interaction | $F_{1,40}$ = 0.07739 | 0.7823 | $F_{1,40}$ = 1.665 | 0.2043 |
| Sex x EtOH Interaction | $F_{1,40}$ = 0.1066 | 0.7457 | $F_{1,40}$ = 0.6681 | 0.4185 |
| Stress x EtOH Interaction | $F_{1,40}$ = 0.1849 | 0.6695 | $F_{1,40}$ = 1.564 | 0.2183 |
| Sex x Stress x EtOH Interaction | $F_{1,40}$ = 0.2382 | 0.6282 | $F_{1,40}$ = 0.2689 | 0.6069 |

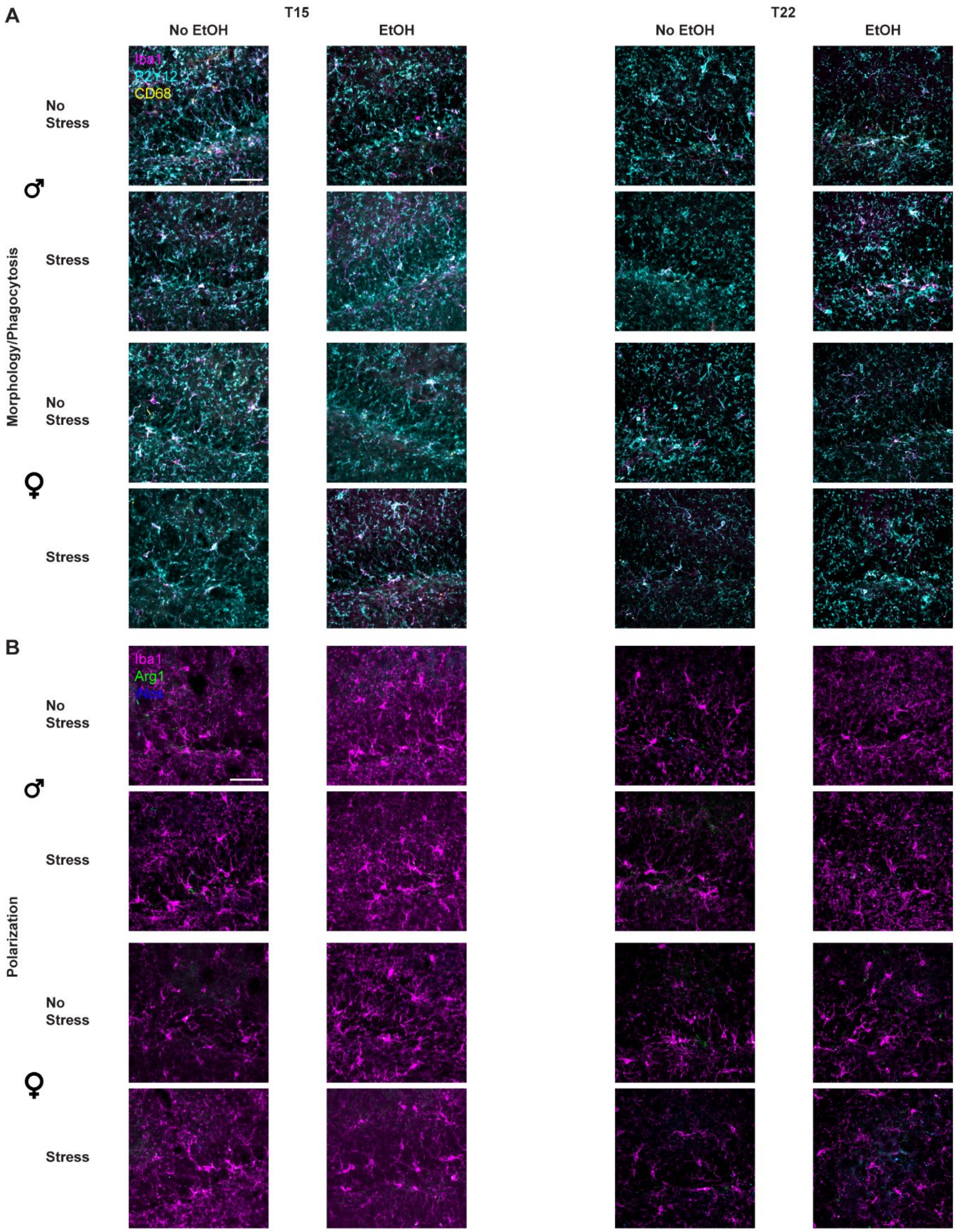

**Fig. S12: Representative micrographs for DG microglia in Experiments 1 & 2, related to Fig. 3.**

All images are shown as Max Intensity Z-Projections. **A**. Representative images for the Morphology/Phagocytosis stain; Iba1 is magenta, CD68 is yellow, and P2Y12 is cyan.

**B**. Representative images for the Polarization stain; Iba1 is magenta, Arg1 is green, and iNos is blue. Scale bar = 50 μm.

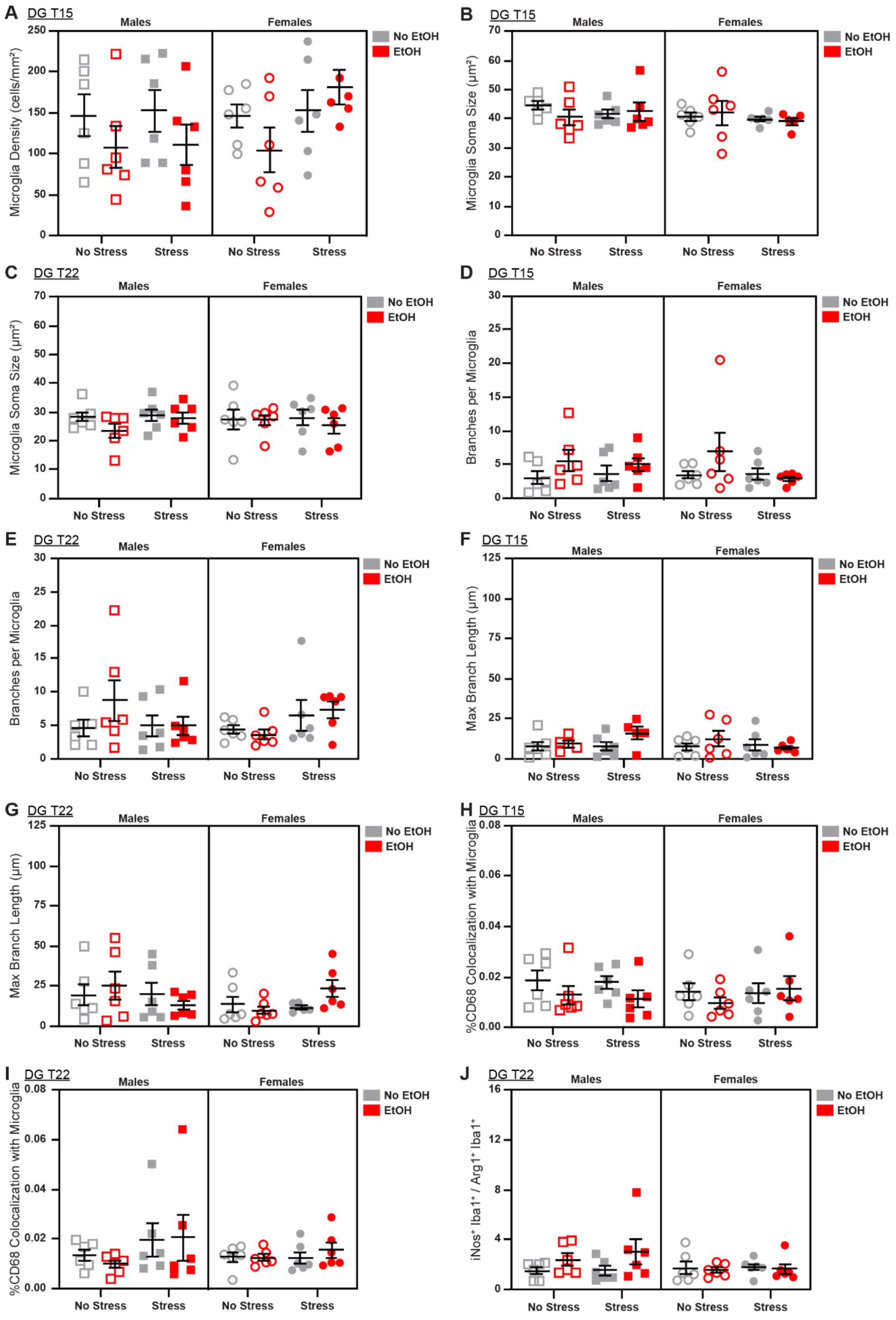

A
DG T15
Males
Females
Microglia Density (cells/mm²)
No Stress
Stress
No EtOH
EtOH
B
DG T15
Microglia Soma Size (µm²)
C
DG T22
Microglia Soma Size (µm²)
D
DG T15
Branches per Microglia
E
DG T22
Branches per Microglia
F
DG T15
Max Branch Length (µm)
G
DG T22
Max Branch Length (µm)
H
DG T15
%CD68 Colocalization with Microglia
I
DG T22
%CD68 Colocalization with Microglia
J
DG T22
iNos⁺ Iba1⁺ / Arg1⁺ Iba1⁺

**Fig. S13: Additional characterization of sex, stress, and EtOH effects on microglial phenotypes in DG from Experiments 1 & 2, related to Fig. 3**
There were no main effects of sex, stress, or EtOH on **A**. microglia density on T15 (1 outlier excluded), **B.** microglia soma size on T15 (2 outliers excluded), **C.** microglia soma size on T22, **D.** branch number on T15, **E.** branch number on T22, **F.** branch length on T15 (2 outliers excluded), **G.** branch length on T22 (1 outlier excluded), **H.** microglial CD68 expression on T15, **I.** CD68 expression on T22, or **J.** M1/M2 polarization on T22 (1 outlier excluded). See Fig. S12 for representative micrographs and Table S6 for detailed statistics. *n* = 5-6 animals/group.

**Table S6: Complete statistical analyses for characterization of DG microglia in Experiments 1 & 2, related to Fig. 3.**
Data shown in Figs. 3 and S13. **p* < 0.05; ***p* < 0.01.

| | T15 | | T22 | |
|---|---|---|---|---|
| **Analysis** | ***F / t*** | ***p*** | ***F / t*** | ***p*** |
| 3-way ANOVA: Microglia Density | | | | |
| Effect of Sex | $F_{1,40}$ = 0.9625 | 0.3324 | $F_{1,40}$ = 0.3730 | 0.5449 |
| Effect of Stress | $F_{1,40}$ = 1.866 | 0.1795 | $F_{1,40}$ = 0.7428 | 0.3939 |
| Effect of EtOH | $F_{1,40}$ = 1.917 | 0.1739 | $F_{1,40}$ = 0.2824 | 0.5981 |
| Sex x Stress Interaction | $F_{1,40}$ = 1.231 | 0.2739 | $F_{1,40}$ = 6.163 | 0.0173* |
| Sex x EtOH Interaction | $F_{1,40}$ = 0.9989 | 0.3236 | $F_{1,40}$ = 0.5324 | 0.4698 |
| Stress x EtOH Interaction | $F_{1,40}$ = 0.9625 | 0.3324 | $F_{1,40}$ = 0.4403 | 0.5108 |
| Sex x Stress x EtOH Interaction | $F_{1,40}$ = 1.191 | 0.2817 | $F_{1,40}$ = 4.187 | 0.0473* |
| 2-way ANOVA: Microglia Density, Males | | | | |
| Effect of Stress | | | $F_{1,20}$ = 1.370 | 0.2555 |
| Effect of EtOH | | | $F_{1,20}$ = 0.8297 | 0.3732 |
| Stress x EtOH Interaction | | | $F_{1,20}$ = 0.9974 | 0.3299 |
| 2-way ANOVA: Microglia Density, Females | | | | |
| Effect of Stress | | | $F_{1,20}$ = 5.369 | 0.0312* |
| Effect of EtOH | | | $F_{1,20}$ = 0.01887 | 0.8921 |
| Stress x EtOH Interaction | | | $F_{1,20}$ = 3.525 | 0.0751 |
| Šídák’s multiple comparisons test: Microglia Density, Females | | | | |

| | | | | |
|---|---|---|---|---|
| No Stress/No EtOH v. No Stress/EtOH | | | $t$ = 1.425 | 0.3105 |
| Stress/No EtOH v. Stress/EtOH | | | $t$ = 1.230 | 0.4114 |
| No EtOH/No Stress v. No EtOH/Stress | | | $t$ = 0.3108 | 0.9420 |
| EtOH/No Stress v. EtOH/Stress | | | $t$ = 2.966 | 0.0152* |
| 3-way ANOVA: Microglia Soma Size | | | | |
| Effect of Sex | $F_{1,39}$ = 1.227 | 0.2748 | $F_{1,40}$ = 0.01689 | 0.8972 |
| Effect of Stress | $F_{1,39}$ = 0.5510 | 0.4623 | $F_{1,40}$ = 0.2511 | 0.6191 |
| Effect of EtOH | $F_{1,39}$ = 0.1288 | 0.7216 | $F_{1,40}$ = 1.634 | 0.2085 |
| Sex x Stress Interaction | $F_{1,39}$ = 0.1554 | 0.6956 | $F_{1,40}$ = 0.8585 | 0.3597 |
| Sex x EtOH Interaction | $F_{1,39}$ = 0.3472 | 0.5591 | $F_{1,40}$ = 0.1253 | 0.7252 |
| Stress x EtOH Interaction | $F_{1,39}$ = 0.2268 | 0.6365 | $F_{1,40}$ = 0.04536 | 0.8324 |
| Sex x Stress x EtOH Interaction | $F_{1,39}$ = 1.188 | 0.2825 | $F_{1,40}$ = 0.8513 | 0.3617 |
| 3-way ANOVA: Branches per Microglia | | | | |
| Effect of Sex | $F_{1,40}$ = 0.01359 | 0.9078 | $F_{1,40}$ = 0.1096 | 0.7423 |
| Effect of Stress | $F_{1,40}$ = 0.9415 | 0.3377 | $F_{1,40}$ = 0.2419 | 0.6255 |
| Effect of EtOH | $F_{1,40}$ = 2.944 | 0.0939 | $F_{1,40}$ = 0.7710 | 0.3851 |
| Sex x Stress Interaction | $F_{1,40}$ = 0.9994 | 0.3235 | $F_{1,40}$ = 3.683 | 0.0621 |
| Sex x EtOH Interaction | $F_{1,40}$ = 0.08172 | 0.7765 | $F_{1,40}$ = 0.6376 | 0.4293 |
| Stress x EtOH Interaction | $F_{1,40}$ = 1.936 | 0.1718 | $F_{1,40}$ = 0.2888 | 0.5940 |
| Sex x Stress x EtOH Interaction | $F_{1,40}$ = 0.5562 | 0.4601 | $F_{1,40}$ = 1.366 | 0.2494 |
| 3-way ANOVA: Max Branch Length | | | | |
| Effect of Sex | $F_{1,38}$ = 0.3227 | 0.5733 | $F_{1,39}$ = 1.357 | 0.2511 |
| Effect of Stress | $F_{1,38}$ = 0.07322 | 0.7882 | $F_{1,39}$ = 2.758e-005 | 0.9958 |
| Effect of EtOH | $F_{1,38}$ = 1.972 | 0.1683 | $F_{1,39}$ = 0.1962 | 0.6603 |
| Sex x Stress Interaction | $F_{1,38}$ = 1.748 | 0.1940 | $F_{1,39}$ = 2.093 | 0.1560 |
| Sex x EtOH Interaction | $F_{1,38}$ = 0.5926 | 0.4462 | $F_{1,39}$ = 0.3419 | 0.5621 |
| Stress x EtOH Interaction | $F_{1,38}$ = 0.0006834 | 0.9793 | $F_{1,39}$ = 0.06454 | 0.8008 |

| | | | | |
|---|---|---|---|---|
| Sex x Stress x EtOH Interaction | $F_{1,38}$ = 2.216 | 0.1499 | $F_{1,39}$ = 2.995 | 0.0914 |
| 3-way ANOVA: %CD68 Colocalization with Microglia | | | | |
| Effect of Sex | $F_{1,40}$ = 0.5531 | 0.4614 | $F_{1,40}$ = 0.7516 | 0.3911 |
| Effect of Stress | $F_{1,40}$ = 0.06827 | 0.7952 | $F_{1,40}$ = 2.476 | 0.1235 |
| Effect of EtOH | $F_{1,40}$ = 2.103 | 0.1548 | $F_{1,40}$ = 9.145e-005 | 0.9924 |
| Sex x Stress Interaction | $F_{1,40}$ = 0.5332 | 0.4695 | $F_{1,40}$ = 1.287 | 0.2634 |
| Sex x EtOH Interaction | $F_{1,40}$ = 1.052 | 0.3112 | $F_{1,40}$ = 0.1927 | 0.6631 |
| Stress x EtOH Interaction | $F_{1,40}$ = 0.3070 | 0.5826 | $F_{1,40}$ = 0.4248 | 0.5183 |
| Sex x Stress x EtOH Interaction | $F_{1,40}$ = 0.5365 | 0.4682 | $F_{1,40}$ = 0.01401 | 0.9064 |
| 3-way ANOVA: $iNos^+$ $Iba1^+$ / $Arg1^+$ $Iba1^+$ | | | | |
| Effect of Sex | $F_{1,40}$ = 0.1617 | 0.6897 | $F_{1,40}$ = 1.581 | 0.2159 |
| Effect of Stress | $F_{1,40}$ = 0.5161 | 0.4767 | $F_{1,40}$ = 0.3160 | 0.5772 |
| Effect of EtOH | $F_{1,40}$ = 0.3480 | 0.5586 | $F_{1,40}$ = 2.157 | 0.1498 |
| Sex x Stress Interaction | $F_{1,40}$ = 1.286 | 0.2636 | $F_{1,40}$ = 0.1133 | 0.7382 |
| Sex x EtOH Interaction | $F_{1,40}$ = 0.8010 | 0.3762 | $F_{1,40}$ = 3.899 | 0.0553 |
| Stress x EtOH Interaction | $F_{1,40}$ = 4.784 | 0.0346* | $F_{1,40}$ = 0.2158 | 0.6448 |
| Sex x Stress x EtOH Interaction | $F_{1,40}$ = 4.422 | 0.0418 | $F_{1,40}$ = 0.1562 | 0.6948 |
| 2-way ANOVA: $iNos^+$ $Iba1^+$ / $Arg1^+$ $Iba1^+$, Males | | | | |
| Effect of Stress | $F_{1,20}$ = 1.413 | 0.2485 | | |
| Effect of EtOH | $F_{1,20}$ = 0.9079 | 0.3520 | | |
| Stress x EtOH Interaction | $F_{1,20}$ = 7.579 | 0.0123* | | |
| Šídák's multiple comparisons test: $iNos^+$ $Iba1^+$ / $Arg1^+$ $Iba1^+$, Males | | | | |
| No Stress/No EtOH v. No Stress/EtOH | $t$ = 1.273 | 0.3879 | | |
| Stress/No EtOH v. Stress/EtOH | $t$ = 2.620 | 0.0325* | | |

| | | |
|---|---|---|
| No EtOH/No Stress v. No EtOH/Stress | $t$ = 2.787 | 0.0226* |
| EtOH/No Stress v. EtOH/Stress | $t$ = 1.106 | 0.4842 |
| 2-way ANOVA: iNos$^+$ Iba1$^+$ / Arg1$^+$ Iba1$^+$, Females | | |
| Effect of Stress | $F_{1,20}$ = 1.099 | 0.7438 |
| Effect of EtOH | $F_{1,20}$ = 0.05922 | 0.8102 |
| Stress x EtOH Interaction | $F_{1,20}$ = 0.004514 | 0.9471 |

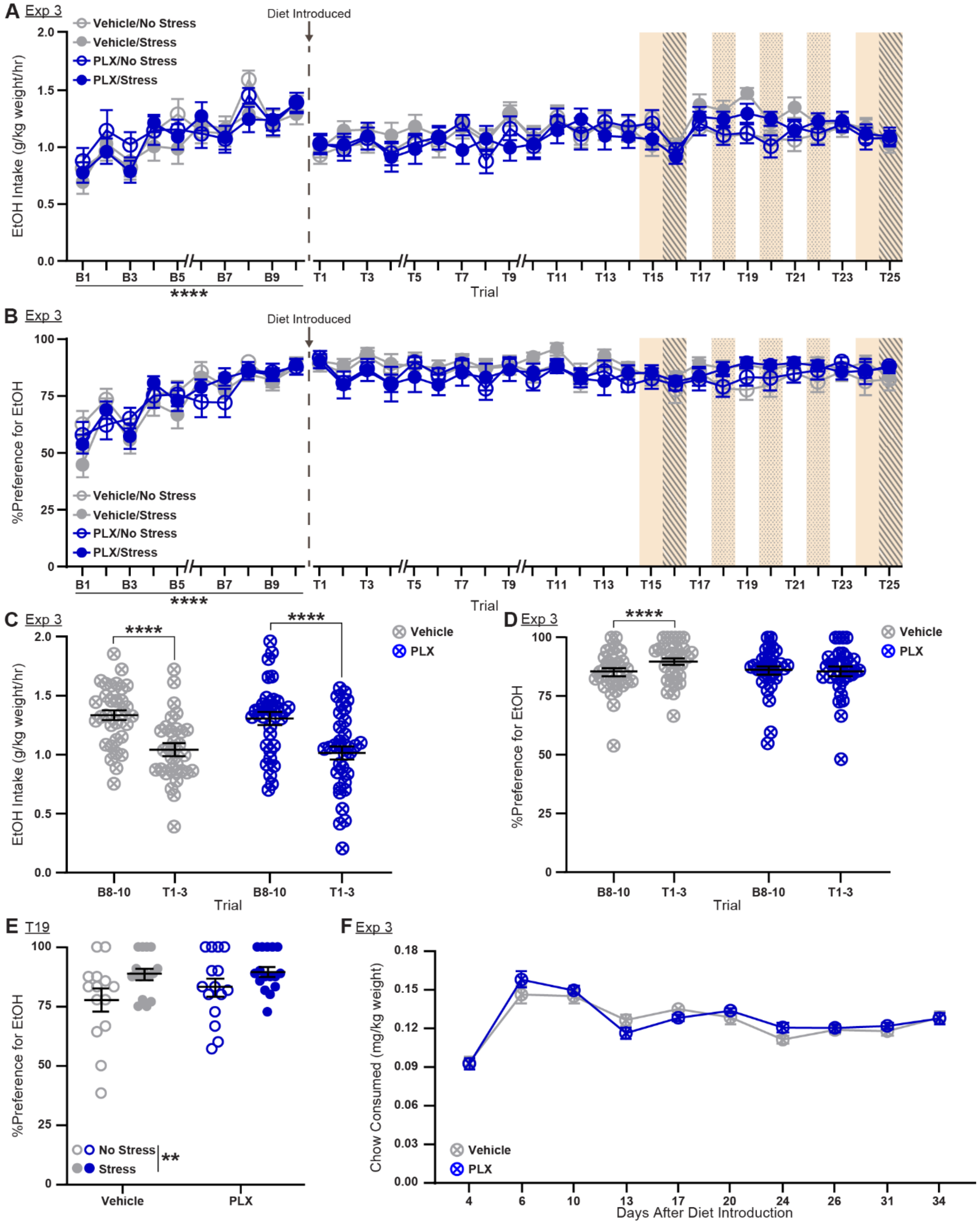

**Fig. S14: Additional characterization of behavioral changes in Experiment 3, related to Fig. 4.**

**A**. Full time course of ethanol intake during 35 trials of DiD (10 pre-diet change, 25 post-diet change) in Experiment 3. Despite an acute reduction in EtOH intake after diet switch, groups did not differ in EtOH intake in the 14 trials after diet introduction. **B**. Full time course of relative preference of EtOH in Experiment 3. In the 14 trials after diet introduction, PLX mice had a slightly lower relative preference for EtOH compared to vehicle mice. **C.** Diet switch caused an acute decrease in EtOH intake across both diet groups, when comparing average EtOH intake in the 3 trials pre- and post-diet change. **D.** Diet switch caused a slight increase in relative EtOH intake, specifically in the vehicle group, but not in the PLX group. **E.** Stress increased relative preference for EtOH on trial 19, an effect prominent in the vehicle group, but not seen in the PLX group. **F.** Mice in the diet and stress subgroups did not differ in chow consumption over the course of 35 calendar days post diet change. $^{**}p < 0.01$; $^{****}p < 0.0001$; $n = 16$/group. Data are averages +/- SEM.

**References for Supplementary Materials**

1. Godynyuk E, Bluitt MN, Tooley JR, Kravitz AV, Creed MC. An Open-Source, Automated Home-Cage Sipper Device for Monitoring Liquid Ingestive Behavior in Rodents. eNeuro. 2019;6.
2. Rhodes JS, Best K, Belknap JK, Finn DA, Crabbe JC. Evaluation of a simple model of ethanol drinking to intoxication in C57BL/6J mice. Physiol Behav. 2005;84:53–63.
3. Abdulla ZI, Mineur YS, Crouse RB, Etherington IM, Yousuf H, Na JJ, et al. Acetylcholine signaling in the medial prefrontal cortex mediates the ability to learn an active avoidance response following learned helplessness training. 2023:2023.09.23.559126.
4. Caldarone BJ, George TP, Zachariou V, Picciotto MR. Gender Differences in Learned Helplessness Behavior Are Influenced by Genetic Background. Pharmacol Biochem Behav. 2000;66:811–817.
5. Caldarone BJ, Harrist A, Cleary MA, Beech RD, King SL, Picciotto MR. High-affinity nicotinic acetylcholine receptors are required for antidepressant effects of amitriptyline on behavior and hippocampal cell proliferation. Biol Psychiatry. 2004;56:657–664.
6. Caldarone BJ, Karthigeyan K, Harrist A, Hunsberger JG, Wittmack E, King SL, et al. Sex differences in response to oral amitriptyline in three animal models of depression in C57BL/6J mice. Psychopharmacology (Berl). 2003;170:94–101.
7. Schindelin J, Arganda-Carreras I, Frise E, Kaynig V, Longair M, Pietzsch T, et al. Fiji: an open-source platform for biological-image analysis. Nat Methods. 2012;9:676–682.
8. Paxinos G, Franklin KBJ. Paxinos and Franklin’s The Mouse Brain in Stereotaxic Coordinates. Compact 5th Edition. San Diego, CA: Elsevier Academic Press; 2019.
9. Young K, Morrison H. Quantifying Microglia Morphology from Photomicrographs of Immunohistochemistry Prepared Tissue Using ImageJ. J Vis Exp JoVE. 2018:57648.
